# Interpretable and explainable predictive machine learning models for data-driven protein engineering

**DOI:** 10.1101/2024.02.18.580860

**Authors:** David Medina-Ortiz, Ashkan Khalifeh, Hoda Anvari-Kazemabad, Mehdi D. Davari

## Abstract

Protein engineering using directed evolution and (semi)rational design has emerged as a powerful strategy for optimizing and enhancing enzymes or proteins with desired properties. Integrating artificial intelligence methods has further enhanced and accelerated protein engineering through predictive models developed in data-driven strategies. However, the lack of explainability and interpretability in these models poses challenges. Explainable Artificial Intelligence addresses the interpretability and explainability of machine learning models, providing transparency and insights into predictive processes. Nonetheless, there is a growing need to incorporate explainable techniques in predicting protein properties in machine learning-assisted protein engineering. This work explores incorporating explainable artificial intelligence in predicting protein properties, emphasizing its role in trustworthiness and interpretability. It assesses different machine learning approaches, introduces diverse explainable methodologies, and proposes strategies for seamless integration, improving trustworthiness. Practical cases demonstrate the explainable model’s effectiveness in identifying DNA binding proteins and optimizing Green Fluorescent Protein brightness. The study highlights the utility of explainable artificial intelligence in advancing computationally assisted protein design, fostering confidence in model reliability.

## 1 Introduction

The advancement of protein engineering, driven by directed evolution (DE) and (semi)rational design (RD), has significantly enhanced protein properties (Wang et al., 2021; Wittmund et al., 2022; Pramanik et al., 2021). These improvements have had far-reaching implications across diverse sectors, including biocatalysis, pharmaceuticals and medicine, livestock, and agriculture (Arnold, 2018). Notably, DE has achieved remarkable success in different applications to optimize and enhance general enzyme properties such as stability and solubility or catalytic properties, e.g., activity and stereoselectivity Pramanik et al. (2021); Arnold (2019). Nevertheless, despite the numerous successful examples, challenges persist, primarily associated with economic constraints, the time-intensive nature of generating and characterizing improved variants, the vast sequence space for exploration, the evaluation of combinatorial variants, and learning epistasis and residue coevolution patterns (Wittmund et al., 2022).

Recently, advancements in deep mutational scanning (DMS), high-throughput screening, and DNA sequencing have accelerated the process of obtaining and analyzing new variants (Vanella et al., 2022). However, the issues of functional classification and the discovery of relevant patterns remain pertinent, underscoring the growing demand for computational methods to support the analysis of evolutionary processes and the exploration of protein functions (Wittmund et al., 2022). The advent of artificial intelligence (AI) and the integration of autonomous systems have ushered in a new era across various domains of scientific research, particularly in studying protein sequences and their multifaceted applications. Notably, the field has witnessed a transformative shift, thanks to the remarkable progress in machine learning (ML) and deep learning (DL) techniques, leading to the development of predictive systems for intricate biological and chemical tasks (Walker et al., 2021).

ML methods have found invaluable applications in diverse biotechnology areas, including drug discovery (Rickerby et al., 2020), assessing various aspects of protein fitness such as thermostability (Csicsery-Ronay et al., 2022), stereoselectivity (Moon et al., 2021; Li et al., 2021), fluorescence properties (Somermeyer et al., 2022), predicting affinity in protein complex interactions (Medina-Ortiz et al., 2023; Liu et al., 2021), functional classification based on Enzyme Commission numbers (Shi et al., 2022; Fernández et al., 2023), recognition of biological activities in peptide sequences (Quiroz et al., 2021), photoreceptor adduct lifetime (Hemmer et al., 2023), and assessing DNA-binding proteins (Qu et al., 2019). The versatility of ML methods has resulted in their integration with traditional experimental techniques, such as DE and RD (Yang et al., 2019; Wittmann et al., 2021). This fusion has given rise to ML-assisted directed evolution (MLDE) and semi-rational design strategies, expanding the toolkit of protein engineering (Kouba et al., 2023; Siedhoff et al., 2020, 2021; Illig et al., 2022). Besides, ML applications extend beyond protein design and have been instrumental in creating predictive models for biotechnological and structural biology challenges, fostering a symbiotic relationship between experimental methods and computational systems (Arnold, 2019).

Despite the increasing success of employing ML methods in data-driven protein engineering, the mathematical models that underlie these advancements often prove challenging for human interpretation (Kouba et al., 2023). Interpretability and explainability are the most common questions for developing predictive models through ML algorithms. Moreover, developing trustworthy predictive models has been one of the most complex challenges to facilitate the application of ML and AI techniques in different medical and biotechnology applications (Kouba et al., 2023). Interpretability in ML refers to understanding and explaining a predictive model’s predictions and decision-making processes comprehensibly to humans (Carvalho et al., 2019). In contrast, explainability refers to understanding how the model learns during the training process (Burkart and Huber, 2021). Explainable Artificial Intelligence (XAI) is a subfield of artificial intelligence that focuses on developing AI systems and ML models that are capable of providing transparent and understandable explanations for their decisions, actions, and predictions (Samek et al., 2019; Holzinger et al., 2022). The primary goal of XAI is to make AI systems more interpretable and understandable to humans, including experts and non-experts (Arrieta et al., 2020). XAI seeks to address the *“black box”* problem commonly associated with complex AI and ML models. Many advanced AI algorithms, such as deep neural networks, can make highly accurate predictions, but their inner workings are often challenging to interpret.

This work addresses the fundamental concepts of Explainable Artificial Intelligence (XAI) and its practical applicability in developing predictive models within the domain of data-driven protein engineering. We delineate the design and implementation of a robust framework for training predictive models tailored to protein engineering tasks, leveraging methodologies from machine learning (ML). Subsequently, we explore XAI’s core characteristics and properties, offering insights into standard methods and their diverse applications. The discussion revolves around innovative strategies for seamlessly integrating XAI methods into developing predictive models specifically designed for protein engineering tasks. To illustrate this integration, we present practical use cases involving classification and regression tasks that combine representation strategies grounded in amino acid encoding through physicochemical approaches, machine learning algorithms, and the application of feature-based and instance-based XAI approaches. These exemplary cases underscore the potential for discerning crucial properties and descriptors relevant to model operation, emphasizing the transformative impact of integrating XAI strategies. Through this integration, we showcase the feasibility of rendering predictive model development transparent, enhancing model explainability, and leveraging acquired knowledge to inform the design of novel protein variants. Lastly, we deliberate on the primary challenges and issues associated with incorporating XAI approaches into conventional data-driven protein engineering models, guiding the effective integration of XAI methods to steer the protein design process.

## 2 Developing predictive models for data-driven protein engineering

Training a predictive model for protein engineering tasks involves generic steps applicable across various model types Kouba et al. (2023); Siedhoff et al. (2020). Figure **1** A illustrates a conventional ML-based predictive model training pipeline employing data-driven techniques. Initially, a dataset comprising the target response variable is gathered, typically related to properties like protein fitness or other desirable characteristics to be predicted and/or enhanced. The main steps required to build an ML sequence–function model and to use those models to guide engineering are introduced below.

**Figure 1:**
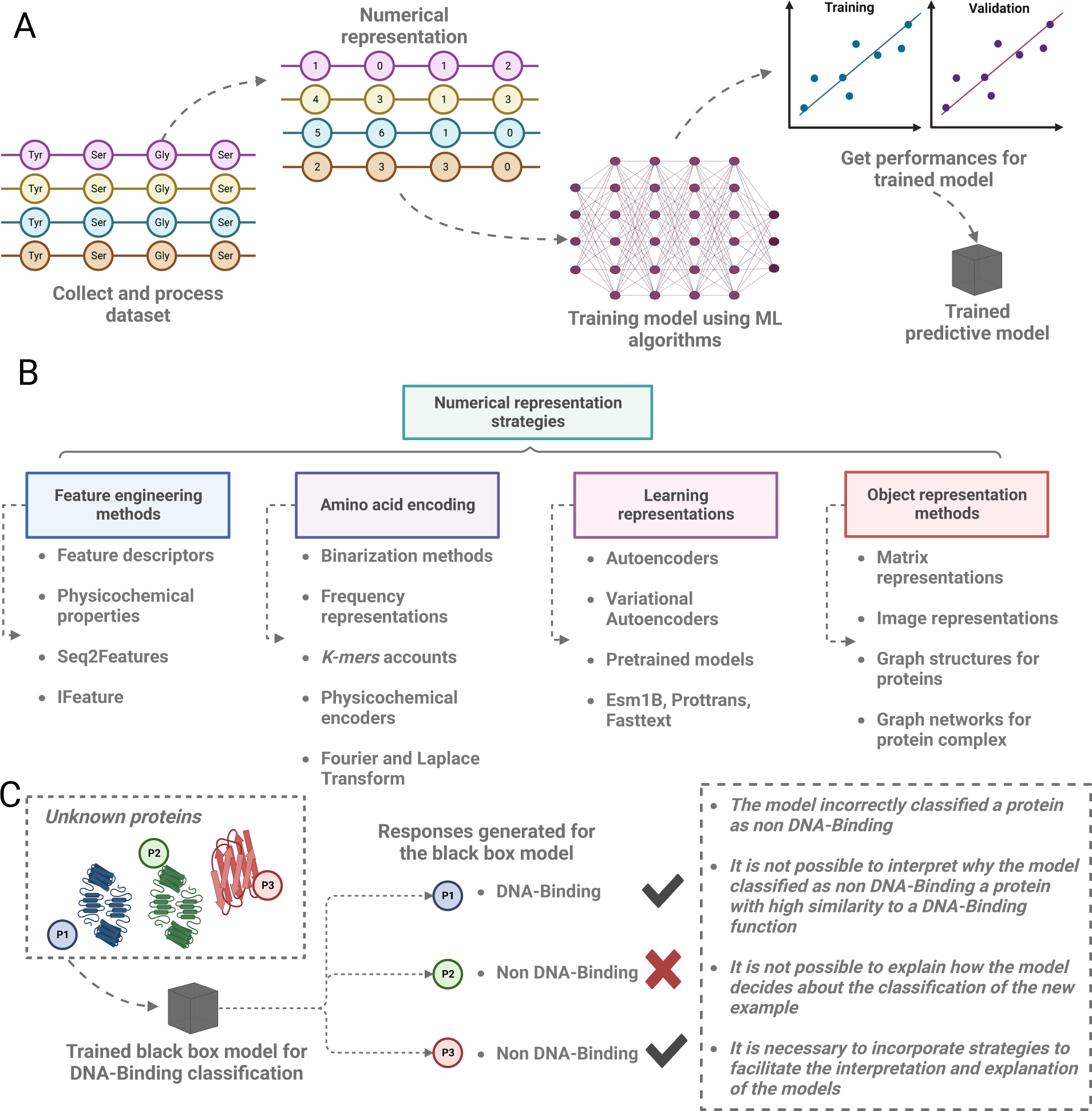
Machine learning strategies applied to build predictive models for protein engineering datasets. **A** A classic pipeline to develop predictive models for protein engineering tasks. First, the protein sequences and the interest response are collected and processed. Then, numerical representation strategies are applied to generate datasets for the training process. Then, an ML algorithm is applied to train a predictive model, and different metrics are employed to evaluate the performance of the trained model. **B** The most common numerical representation strategies applied for protein engineering include feature engineering methods, amino acid encoding approaches, learning representations, and object representation methods. **C** A DNA interaction protein classification black box model is exemplified to demonstrate the consequences of the design and implementation of black box models. We work with unknown proteins and use the model to classify them. During the validation process, two proteins are classified correctly. However, a protein is identified incorrectly, making it complex to interpret why the model does so. This demonstrates the need for XAI strategies to facilitate the interpretation and explanation of the predictions generated by predictive models.

### 2.1 Data collection and processing

Datasets are compiled from various sources, encompassing repositories, literature databases, experimentally generated recombinant libraries, and data originating from specialized experiments such as deep mutational scanning Mazurenko et al. (2020). Notably, the MaveDB database (Rubin et al., 2021) stands out as a valuable resource, hosting an extensive collection of over 100 deep mutational scanning experiments. This includes investigations involving essential proteins like Angiotensin-converting enzyme 2 (ACE2) (Chan et al., 2020), activation-induced deaminase (AID) (Gajula et al., 2014), SUMO1 (small ubiquitin-like modifier), and TPK1 (thiamin pyrophosphokinase) (Weile et al., 2017).

Following the compilation of datasets, various strategies are applied to process and refine the data. These strategies involve the removal of noise or outlier elements, as well as the elimination of redundancy or duplication, among other cleaning procedures. This meticulous cleaning process is crucial for mitigating biases during the training of predictive models, fostering a more robust and generalized predictive model performance (Medina-Ortiz et al., 2020). Moreover, the overall data refinement process can be improved by leveraging imputation techniques to enhance completeness and coherence. Statistically relevant data can be synthesized, or the dataset can be normalized to correct missing values (Song and Shepperd, 2007).

### 2.2 Sequence representation

After acquiring the datasets and their associated response variables, the data must be processed and represented numerically for interpretation by ML algorithms. Figure **1** B overviews the most common numerical representation techniques, including feature engineering and learning representations through pre-trained models.

Among these techniques, feature engineering methods utilize protein descriptors to represent protein sequences based on their properties. These descriptors encompass physicochemical, thermodynamic, evolutionary, and structural attributes. Tools like Seq2feature (Nikam and Gromiha, 2019), ProFET (Ofer and Linial, 2015), and FEGS (Mu et al., 2021) are frequently employed to derive these descriptors. Feature engineering also includes strategies based on amino acid composition (AAC). These methods use the frequency of amino acids in a protein sequence as description methods and incorporate physicochemical properties to describe the residues fully (Ismail et al., 2017). Alternatively, amino acid coding methods generate numerical vectors encoding individual amino acids. These methods include one-hot encoding, ordinal encoding, physicochemical property-based encoding, frequency-based encoding (Siedhoff et al., 2020; Medina-Ortiz et al., 2022), and DCA-based encoding Illig et al. (2022). Ensuring uniform vector sizes may require techniques such as *zero-padding*. Siedhoff et al. (2020), Medina-Ortiz et al. (2022), and Cadet et al. (2018) have even proposed spatial transformations post-encoding, using Fourier or Laplace transforms based on physicochemical properties.

A noteworthy strategy is the use of *k-mers*-based encoding, famous for its applicability in Natural Language Processing (NLP) methods due to its ability to capture context and semantics of protein segments (Guo et al., 2021). Recently, the pivotal role of designing and implementing representation learning models has come to the forefront in developing numerical representation methods for proteins (Ofer et al., 2021). These models draw inspiration from strategies employed in large language models and leverage architectures like autoencoders, variational autoencoders, and similar frameworks to construct sophisticated representation learning systems (Thirunavukarasu et al., 2023). Notable tools such as EcNet (Luo et al., 2021) and TAPE (Rao et al., 2019) have been devised to streamline the numerical representation of proteins and introduce the application of transfer learning through fine-tuning methods. Additionally, specialized libraries like bio-embedding have emerged to simplify using diverse pre-trained models, including UniRep, Bepler, Esm1B, Prottrans, and Seq2vec (Dallago et al., 2021). These advancements collectively contribute to a more nuanced and effective approach in the dynamic landscape of protein representation methods.

From a structural perspective, proteins can also be represented using matrix strategies that combine physicochemical properties with three-dimensional protein data to construct visual representations. Additionally, graph-based techniques have been implemented to connect information and represent atoms, residues, and interactions. Finally, several other encoding methods for protein representation exist, including graph structure representation learning (Yang et al., 2020), geometric graph structure representation learning (Xia and Ku, 2021), weighted sparse representation matrices (Huang et al., 2016), and chaos-game-based representations (Sun et al., 2020).

### 2.3 Predictive ML model training and evaluation

Once the datasets undergo numerical representation, the training of predictive models proceeds using either ML algorithms or DL architectures. The initial step involves partitioning the dataset into training and validation data sets, usually split using a classic proportion of 70:30 or 80:20. The training data set is utilized to train and optimize the predictive model’s hyperparameters. In contrast, the validation data set assesses the model’s performance (Medina-Ortiz et al., 2020). A range of supervised learning algorithms can be employed, such as Random Forest, Support Vector Machine, and DL architectures like Recurrent Neural Networks (RNN), Long Short-Term Memory (LSTM), and Convolutional Neural Networks (CNN) Kouba et al. (2023); Mazurenko et al. (2020). Evaluation metrics, tailored to the model type, include precision, recall, Matthews Correlation Coefficient (MCC) for classification, and Pearson coefficient, mean square error for regression models. Strategies like *k*−fold cross-validation are applied to prevent the overfitting. Alternatively, applying test datasets could generate the same effect as cross-validation methods. Finally, a tuning hyperparameter process is usually applied to optimize the model’s hyperparameters.

Implementing an ML-based predictive model for protein engineering tasks is indeed viable. However, several crucial factors demand consideration during model development. These encompass data dimensionality, data representation methods, response variability or class imbalance, training strategies, explored algorithms, overfitting assessment, and the replicability of predictive models. Addressing these considerations ensures a robust and practical application of ML in protein engineering.

### 2.4 Interpretability and explainability: The main challenges of the development of predictive models

Constructing predictive models necessitates ensuring confidence among users who employ the models and rely on the responses they generate (Doran et al., 2017). In protein engineering, predictive models are designed to facilitate the creation of proteins with desirable properties (Wittmann et al., 2021). Their primary focus includes classifying proteins with unknown functions, analyzing mutational variants, and aiding in reconstructing landscapes (Wittmund et al., 2022). Consequently, their application revolves around generating predictions crucial for experimental settings (Wittmann et al., 2021). The reliability of a model becomes paramount as an unreliable one could introduce uncertainty during experimental validation, potentially resulting in economic losses should inconsistencies arise between generated predictions and experimentally obtained values (Gale et al., 2019). These challenges in reliability prompt questions about the operational aspects of predictive models—specifically, the *“how”* and *“why”*. These inquiries center around the twin pillars of interpretability and explainability Samek et al. (2019).

A practical example is presented in Figure **1** C. This scenario uses a *black box* classification model for DNA-binding proteins to evaluate three proteins with unknown functions. Among them, proteins *P*_1_ and *P*_2_ represent mutational variants with a point substitution. The model classifies protein *P*_1_ as DNA-Binding, while proteins *P*_2_ and *P*_3_ are categorized as non-DNA-binding proteins. The results indicate the model’s intuitive behavior for proteins *P*_1_ and *P*_3_, as they exhibit apparent structural differences. Protein *P*_1_ aligns with the characteristics of a DNA-binding protein, whereas protein *P*_3_ notably lacks these traits. However, protein *P*_2_ presents a challenge by not conforming to intuitive expectations. Ideally, the model should classify *P*_2_ as a DNA-binding protein but categorize it differently without DNA interaction. This divergence highlights an interpretability challenge in the model’s functionality. Simultaneously, the lack of comprehension regarding how the model generates predictions raises concerns about explainability. Both interpretability and explainability play pivotal roles when employing predictive models, providing the necessary support for model predictions and justifying their actions. These concepts are integral components of explainable artificial intelligence Holzinger et al. (2022).

## 3 Explainable artificial intelligence (XAI): Definitions and characteristics

Explainable Artificial Intelligence (XAI) is a concept focused on the development of artificial intelligence systems and algorithms that empower humans to comprehend, interpret, and have confidence in the decisions and outcomes generated by these systems (Burkart and Huber, 2021; Samek et al., 2019). The primary goal of XAI is to improve the transparency and comprehensibility of AI systems, especially in scenarios where the decision-making processes of AI can significantly impact individuals or organizations (Arrieta et al., 2020; Holzinger et al., 2022). XAI often intersects with interpretable AI or Explainable Machine Learning (XML). It encompasses AI systems that enable humans to maintain intellectual oversight or the techniques employed to facilitate such oversight (Vilone and Longo, 2021). XAI acts as a countermeasure to the *“black box”* phenomenon in machine learning, where the designers of the AI may struggle to articulate the reasons behind a specific decision (Castelvecchi, 2016).

XAI encompasses several essential elements, including i) Transparency: XAI systems are intentionally designed to be transparent, offering insights into the rationale behind their specific conclusions and decisions. This transparency empowers users, stakeholders, and regulatory authorities to understand the underlying logic behind AI recommendations. ii) Interpretability: The concept of interpretability in XAI centers on making AI models and their inner workings more accessible and understandable to humans. This may involve providing clear and intuitive visualizations, explanations, or other methods to convey the reasoning behind AI decisions. iii) Accountability: XAI strongly emphasizes ensuring that AI systems can be held accountable for their actions. When AI decides, a clear trail should lead back to specific data, features, or model parameters, making identifying and addressing errors or biases simpler. iv) Fairness: Ensuring fairness within AI systems is a critical component of XAI. This involves preventing and mitigating biases in AI models to avoid discrimination and unequal treatment of different groups of individuals. v) Trustworthiness: XAI establishes trust between humans and AI systems. Users who comprehend and have confidence in AI decision-making are more inclined to embrace and apply these technologies across various applications (Vilone and Longo, 2021; Samek et al., 2019; Holzinger et al., 2022).

Different strategies exist for developing transparent predictive models through applying XAI methods. As illustrated in Figure **2**, a comprehensive overview of the top five XAI methods is presented (Jiménez-Luna et al., 2020). These approaches are centered around i) determining the relevance of features based on predictions (feature attribution), ii) identifying essential data subsets crucial for accurate model predictions (instance-based), iii) employing methods that interpret information flow in models through convolution strategies (graph convolution-based), iv) incorporating predictive methods designed to be inherently explanatory (self-explaining), and v) employing methods to assess the reliability of predictions made by the generated models (uncertainty estimation). The main methods are described below, along with their main characteristics, development strategies, and different challenges related to the application of the methods.

**Figure 2:**
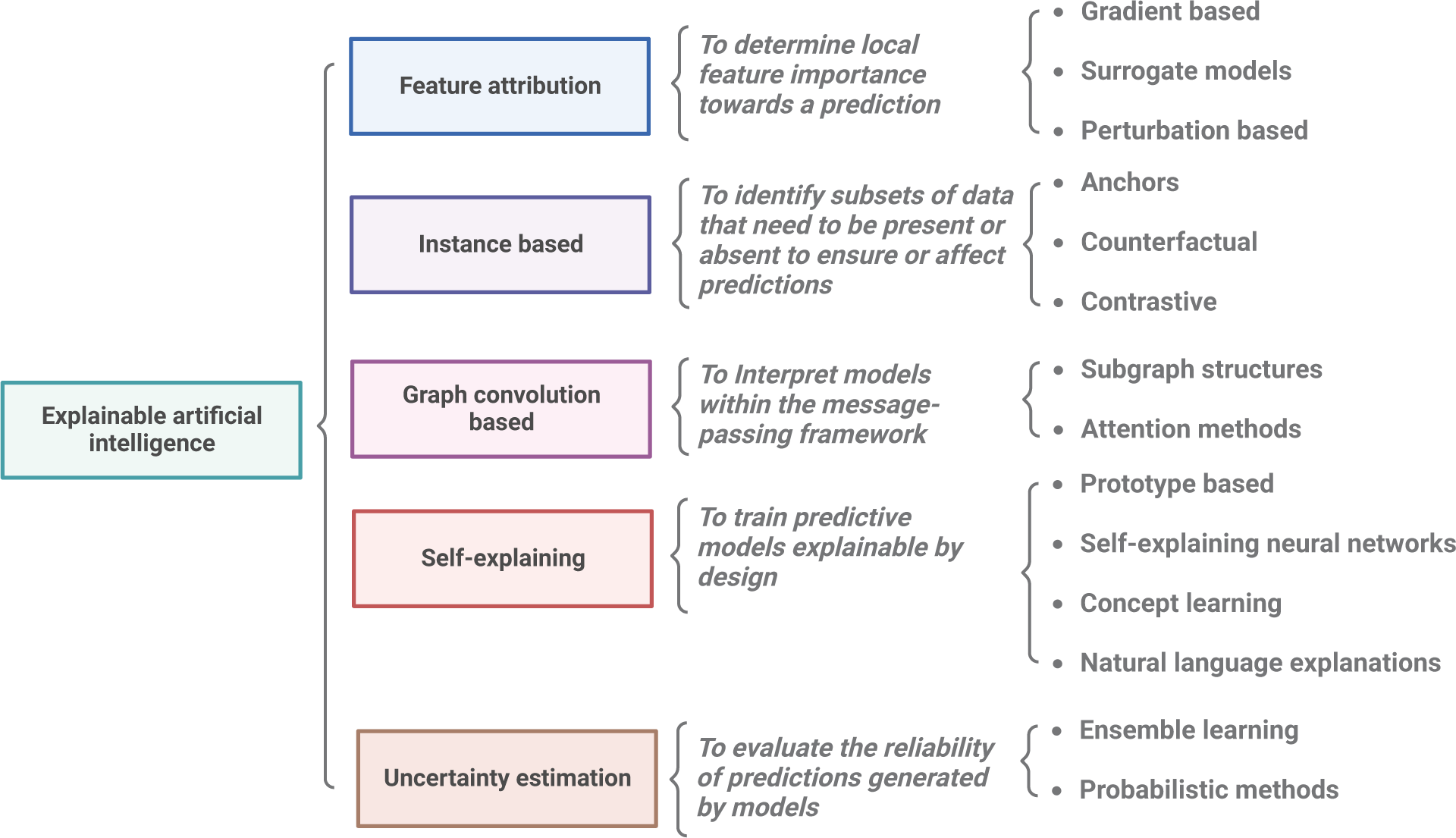
Main explainable artificial intelligence (XAI) methods and strategies with possible applications into protein engineering problems. Explainable Artificial Intelligence (XAI) strategies can be systematically categorized based on their distinctive properties and characteristics. Feature attribution methods, for instance, explore property relevance by analyzing predictive responses or employing gradient-based approaches within neural network architectures. Conversely, instance-based techniques pinpoint crucial areas or attributes influencing model predictions. This category encompasses methods like anchors, counterfactuals, and contrastive approaches. Leveraging graph representations introduces another avenue for XAI. Techniques rooted in subgraph identification or attention methods allow for the comprehensive analysis of nodes, facilitating the identification of relationships with predictive outcomes. Finally, self-explaining and uncertainty are integral components of an XAI strategy. These encompass the development of prototypes or self-explaining neural networks, as well as utilizing ensemble learning and probabilistic methods. This comprehensive framework provides a nuanced understanding of the interpretability and transparency of artificial intelligence models, contributing to their broader adoption and trustworthiness.

### 3.1 Feature attribution methods

Feature attribution methods play a pivotal role in enhancing the transparency and interpretability of ML models by offering insights into the impact of individual input features on predictions. These methods are categorized into local and global models, focusing on explaining individual or all samples, and model-centric and data-centric models, which operate on specific models or measure feature informativeness across the dataset (Gevaert et al., 2022). This work primarily focuses on the main category, which includes gradientbased, surrogate-model, and perturbation-based methods (Jiménez-Luna et al., 2020). These approaches unveil feature importance by introducing changes around the feature’s locality, adding noise to the intended feature, and making deletions or alterations to the feature (Gevaert et al., 2022; Jiménez-Luna et al., 2020). Figure **2** shows the three main approaches of feature attribution methods that could be applied in protein engineering predictive models. These approaches are explained below.

Surrogate models are simplified and interpretable models trained to approximate the behavior of complex, often *“black box”* ML models. The generation of surrogate models involves subsampling instances from the input dataset, making predictions using the *“black box”* model, and training a more interpretable model using the generated subsampling. This interpretable model, such as regression, decision trees, or rule-based methods, clearly represents the complex model’s decision-making process. Interpreting predictions from the surrogate model is generally more straightforward than directly interpreting the complex model. While surrogate models confront the challenge of replicating predictions efficiently from the black-box model, a key question arises: does achieving clear and efficient replication unnecessarily render the need for a complex algorithm to generate predictions on the original dataset? Additionally, the complexity involved in training surrogate models is notable, often employing methods such as decision trees or linear regression, albeit without a guaranteed correct parameter fit. The careful selection of data for developing the surrogate model emerges as a crucial factor, focusing on instances that significantly impact model interpretability and, consequently, explainability.

Gradient-based approaches leverage information from a model’s gradients (derivatives) concerning its inputs to understand how changes in input features impact predictions. These approaches include gradient attribution, integrated gradients, layer-wise relevance promotion (LRP), and guided backpropagation. Gradientbased approaches are particularly valuable for interpreting complex models like deep neural networks. Analyzing the gradients can identify features or regions in the input space that influence predictions, contributing to transparency and interpretability. Combining predictive models trained through DL architectures with gradient-based methods facilitates the explanation of predictions by assessing the impact caused by changes in descriptors. While these strategies enhance the explainability of a deep learning model, the number of nodes, layers, and types of layers during data processing and learning incurs a significant computational cost that scales as layers and nodes expand. Additionally, similar to perturbation-based methods, the application of clustering or pattern identification methods is required to generalize the rules that enable the explanation of predictions generated by the model, adding extra complexity to the training of predictive models.

Perturbation-based approaches involve making controlled changes to input data to observe corresponding changes in the model’s predictions. Methods like LIME, SHAP, and Randomized What-If Analysis are focused on perturbation-based approaches. These techniques help understand a model’s sensitivity to variations in input features, providing valuable insights into feature importance. Perturbation-based approaches benefit complex models, offering actionable insights into decision boundaries and highlighting the contribution of individual features. This contributes to the interpretability of machine learning models by elucidating how changes in input data influence predictions (Jiménez-Luna et al., 2020). Explanation methods based on perturbations, such as LIME or SHAP, appear particularly valuable as they identify relevant variables for explaining a model-generated response. Applying these analyses to all relevant examples underscores the necessity of generalizing these rules. These analyses prove insightful in datasets where the number of descriptors is not high, signifying low dimensionality and a diminished tendency for high nonlinearity in the datasets. However, in datasets with multiple descriptors, the role of these strategies transforms the explanation challenge into a complex data science problem. Generalizing descriptor behaviors into the identified rules for model predictions demands intricate clustering strategies, amplifying the complexity of the problem.

### 3.2 Instance-based approaches

The Instance-Based or instance-based learning (IBL) algorithm differs from traditional methods by evaluating the target function uniquely for each new instance rather than globally for the entire instance space. It leverages classification and regression techniques such as nearest neighbor, case-based reasoning, and locally weighted regression (Aha et al., 1991). Noteworthy for its distinctive feature of putting off generalization until a new instance needs classification, this method is called “lazy” or memory-based learning. To enhance real-world applicability and maximize the classification accuracy of new cases, IBL, exemplified by the *k*-nearest neighbor (IBK), benefits from voting and instance weights through the Distance-weighted Nearest Neighbor Algorithm (Jiménez-Luna et al., 2020; Kanwal and Bostanci, 2016). Praised for its simplicity, efficient time utilization, and ability to maintain high accuracy by favoring specific instances over pre-compiled abstractions in prediction tasks (Kanwal and Bostanci, 2016; Aha et al., 1991), IBL approaches can be categorized into anchor algorithms, counterfactual instance search, and contrastive explanation methods (Jiménez-Luna et al., 2020). These methods are elucidated below.

Anchor explanations represent a specialized approach designed to furnish interpretable justifications for machine learning model predictions, especially in classification tasks (Tan et al., 2023). This method offers concise, understandable, and semantically meaningful statements describing a high-confidence region around a prediction. As if-then rules, anchors articulate the input space conditions strongly correlating with a specific model prediction. The generation process for anchor methods encompasses various approaches, including data perturbation, model querying, and rule construction. Anchor methods prioritize a precision-recall trade-off, seeking rules that balance precision and recall while maintaining human-readable outputs for user comprehension (Mirzaei et al., 2023).

Counterfactual approaches involve generating alternative instances or scenarios to illustrate potential outcomes if specific features or conditions were altered, resulting in a different model prediction (Stepin et al., 2021). Counterfactual instances unveil insights into an ML model’s decision-making process. Counterfactuals are hypothetical instances resembling the original input data but differing in specific features. Users gain insights into feature importance and the conditions leading to precise predictions by observing how the model’s prediction changes with modified features (Dandl et al., 2020). Like anchor methods, counterfactual approaches entail a generation process employing perturbation, model prediction, and explanation methods. Counterfactuals find applications in generating personalized explanations for individual predictions and assessing model fairness by exploring instances where changing sensitive attributes significantly impacts predictions (de Oliveira and Martens, 2021).

Contrastive approaches focus on elucidating ML model predictions by contrasting them with alternative scenarios or instances. The primary objective is to underscore the distinctive factors that contribute to the predicted outcome compared to other conceivable outcomes (You et al., 2020). Contrastive explanations involve a comparative analysis of the prediction for a specific instance with the predictions that would have arisen under varied conditions or scenarios. These scenarios may include introducing changes to the input features while maintaining other aspects constant or utilizing predictions generated by the *“black box”* model in both the original and contrastive scenarios. The spectrum of contrastive approaches includes individual- and group-level contrasts, offering flexibility in the depth of analysis. Furthermore, contrastive approaches can serve both individual explanations and model sensitivity analysis (Zimmermann et al., 2021).

Both anchor and counterfactual methods aim to enhance explainability by constructing simple and descriptive rules that shed light on predictions made by *“black box”*. However, they diverge in their rule structures; anchor explains predictions through if-then conditions, while counterfactual focuses on understanding what changes would lead to a different outcome. The latter allows for simulating behaviors that influence a shift in the model’s response, aiding in comprehending the sensitivity of the response to such alterations.

In contrast, explanation methods based on contrastive approaches share similarities with anchor and counter-factual methods. Applying these methods incurs computational costs, which do not necessarily scale directly with the dataset’s dimensionality. Nevertheless, anchor-based explanatory methods generally involve lower complexity and computational costs than counterfactual and contrastive methods. Besides, counterfactual and contrastive methods introduce extra complexity from a computational standpoint and in interpreting changes made to the descriptors that result in alterations to the model’s response. This interpretation is case-dependent and may often lead to aberrant changes, necessitating validation through expert criteria.

In all instance-based cases, applying sophisticated computational methods focused on pattern identification is essential to facilitate the generalization of rules generated by the widespread use of anchor or counterfactual methods. This introduces additional complexities to the task of efficiently training predictive models.

### 3.3 Graph convolution-based methods

Graph Convolutional Neural Networks (GCNs) have gained widespread popularity as a means of learning graph representations for tasks involving structured data, such as molecular biology, social networks, semantic segmentation, and image classification (Zhang et al., 2018; Wu et al., 2019; Veličkovič et al., 2018). The utility of graph representation learning extends to addressing challenges in various domains, including computer vision, traffic prediction, and natural language processing, by employing embedding techniques in low-dimensional spaces (Li et al., 2022; Chen et al., 2020).

Within Explainable Artificial Intelligence (XAI), a prominent method harnessing GCNs includes subgraph identification and attention-based approaches. Subgraph identification methods aim to maximize information from the relationship between a subgraph structure and a small subset of node features concerning the predicted graph. This formulation transforms the problem into an optimization challenge (Jiménez-Luna et al., 2020). Examining which subgraphs are pivotal for specific model predictions can enhance interpretability. Analyzing subgraph structures involves identifying highly influential features or feature combinations within subgraphs that significantly contribute to the model’s output (Ying et al., 2019).

Adopting the attention mechanism from Natural Language Processing (NLP), attention-based approaches determine hidden node-level representations by stacking multiple message-passing layers. This involves computing an attention coefficient for each edge over the neighbors of its nodes (Jiménez-Luna et al., 2020). These approaches empower models to focus on specific parts of input data during predictions. Attention mechanisms, commonplace in neural networks, enhance model interpretability by emphasizing the importance of different input features. They assign varying weights or importance scores to input data elements, indicating their relevance to the model’s prediction. The dynamic adjustment of these weights during model computation is often visualized as a crucial aspect of attention methods, encompassing self-attention, soft attention, and intricate attention (Veličkovič et al., 2018).

Explanations based on graph methods present a specific approach for elucidating predictive models built on Graph Convolutional Network (GCN) architectures. Both the utilization of subgraphs and the incorporation of attention layers facilitate the explanation of predictive models relying on these architectures. However, the computational cost of identifying and processing subgraphs becomes significant as the dataset’s dimensionality increases. Furthermore, distinct considerations must be explored regarding the application of graph architectures, as they hinge on the nature of the prediction to be developed.

Beyond computational costs, it is evident that post-processing techniques or pattern identification are necessary to foster the generalization of rules that explain model predictions. Additionally, an evaluation based on expert criteria is imperative to interpret the identified patterns. Identifying subgraphs or correlating attention layers merely designates points of interest, which may not inherently provide descriptive insights for interpreting or explaining the model.

In this context, combining subgraph analysis with attention methods demands discrete mathematical modeling techniques to generate more descriptive rules for human understanding. This introduces complexity in computational costs and necessitates expert judgment and additional techniques, posing additional challenges to predictive modeling via GCN and its explanation through graph-based methods.

### 3.4 Self-explaining approaches

To tackle the challenge of interpretability, the concept of self-explaining AI models has emerged, aiming to provide predictions and understandable explanations. Self-explaining integrates transparency by embedding explanations and interpretations into its core design, bolstering trust through mutual information metrics and offering confidence levels for decisions and explanations (Grange et al., 2022; Jiménez-Luna et al., 2020). In a broader context, self-explaining approaches strive to demystify complex systems by incorporating human-interpretable explanations, fostering transparency, trust, and improved decision-making across various applications (Elton, 2020). Several approaches fall under self-explaining methods, including prototype-based methods, self-explaining neural networks, concept learning, and natural language explanations, each detailed below.

Prototype-based approaches involve the creation of representative prototypes capturing the characteristics of different classes or groups in a dataset. These prototypes serve as interpretable data representatives by encapsulating the essential features of various classes explaining model predictions. There are two strategies for generating prototypes, including centroid prototypes, based on computing centroid points for specific classes, and exemplary prototypes, selected as highly representative points per class. These approaches contribute to model transparency and interpretability by furnishing users with concrete examples that elucidate how the model distinguishes between different classes, particularly beneficial when users seek intuitive and tangible explanations for model predictions (Kim et al., 2014; Li et al., 2017).

Self-explaining Neural Networks (SENNs) represent a distinct class of neural networks with intrinsic mechanisms for generating explanations and predictions (Norrenbrock et al., 2023). SENNs aim to enhance transparency and interpretability in complex ML systems. Usually comprising two main components—a prediction network for making predictions and an explanation network for generating human-readable justifications—SENNs provide attention maps or feature importance scores highlighting influential parts of the input (Park and Hwang, 2023). Trained concurrently to optimize accurate predictions and meaningful explanations, SENNs find applications in crucial domains such as healthcare, finance, and natural language processing (Ren et al., 2023).

Concept learning involves extracting and understanding abstract concepts or patterns from data. It transcends memorizing specific instances to recognize and represent higher-level concepts effectively. XAI emphasizes making these learned concepts interpretable and explainable, understanding how the model categorizes instances based on abstract patterns. Concept learning balances complexity and interpretability, recognizing that more straightforward concepts may be more interpretable but might not capture the full nuances of complex data (Jiménez-Luna et al., 2020).

Natural language explanations entail providing human-understandable descriptions to elucidate the decisions or predictions made by machine learning models. These explanations, expressed in natural language, bridge the gap between complex model internals and user understanding, leveraging interpretability techniques such as feature importance, attention mechanisms, or rule extraction. Tailored to the intended audience’s expertise, these explanations empower users to make informed decisions based on the model’s output (Lai et al., 2023).

In self-explaining strategies, both prototype and concept learning methods focus on identifying key patterns to define or characterize the responses generated by predictive models. While this pattern recognition allows describing specific classes or ranges of values, it does not necessarily provide an easily interpretable human understanding and a clear explanation of predictions. This limitation stems from its reliance on the features or information considered during the predictive model training. For instance, in tasks like image classification, explanations typically involve the identification of key pixels.

In methods that integrate various forms of representation and preprocessing, utilizing techniques such as PCA, transformations, etc., explanations are constrained to patterns derived from already transformed values. This necessitates strategies that can effectively describe the initial values of the descriptors employed. Consequently, as the complexity of the initial representation or characterization of examples used for model development increases, the interpretation becomes more challenging for self-explaining strategies, requiring additional methods and stages to achieve clarity.

This complexity also extends to methods based on natural-language explanations and Self-Explaining Neural Networks (SENNs), both relying on generating explanations during model training. However, their efficacy is influenced by the chosen characterization or representation strategy, and they introduce increased computational costs by incorporating processing layers and rule generation. These rules, in turn, demand human interpretation and expert judgment for meaningful processing.

### 3.5 Uncertainty estimation

Numerous endeavors have been undertaken to enhance the uncertainty estimation of deep neural network learning models, a critical aspect for improving their interpretability and reliability in practical applications (Deng et al., 2023). Quantifying prediction errors is integral for making more informed and consistent decisions, particularly in medicinal chemistry and optimization processes. This quantification can be obtained through various sources, including entropy metrics, average probabilities, and variance (Jiménez-Luna et al., 2020; Arkov, 2022). Two significant approaches to uncertainty estimation are ensemble methods and probabilistic learning approaches (Jiménez-Luna et al., 2020).

Ensemble approaches boost neural network models’ accuracy and predictive performance by combining identical models with different input features. This combination mitigates biases in predictions (Jiménez-Luna et al., 2020). Ensemble learning aims to enhance overall predictive performance and interpretability using multiple machine learning models known as an ensemble. These methods combine individual model predictions to improve accuracy and robustness, often providing more transparent insights into the decision-making process. Bagging, boosting, and stacking are common types of ensemble methods. While ensemble methods elevate predictive performance, they may pose challenges in interpretability. The combination of predictions from multiple models can obscure the specific factors influencing a particular decision. Typically, techniques like feature importance, attention mechanisms, and post hoc interpretability are applied as XAI methods to elucidate predictions and interpret complex *“black box”* models (Hansen and Salamon, 1990).

Probabilistic methods utilize probability distributions to model uncertainty, offering nuanced insights into the decision-making process of machine learning models. These methods enhance transparency by expressing predictions in terms of probabilities, conveying the confidence or uncertainty associated with each prediction. Bayesian methods, a common probabilistic approach, involve modeling uncertainty using Bayes’ theorem, updating beliefs based on evidence, and expressing predictions as probability distributions. In this approach, neural network parameters are assigned a prior distribution, and a posterior distribution is computed based on training data to quantify predictive uncertainty (Lakshminarayanan et al., 2017). Probabilistic methods facilitate explanations by allowing practitioners to communicate uncertainty to end-users. Instead of providing a deterministic answer, the model expresses a range of possible outcomes and their associated probabilities. While probabilistic methods offer richer insights, they can pose challenges regarding computational complexity and the need for sophisticated inference techniques. Nonetheless, probabilistic methods in XAI contribute to a more nuanced and interpretable understanding of machine learning model predictions. By explicitly modeling uncertainty, these methods enable practitioners and end-users to make more informed decisions and better comprehend the reliability of the model’s outputs.

Explanations derived from ensemble methods involve concurrently using distinct predictive models employing different descriptors. The synergy from applying these methods enhances the overall model performance. Besides, feature analyses can be employed to pinpoint descriptors that wield a pivotal influence on predictions. However, despite their utility, ensemble methods do not clearly understand the effects of permutations or changes in descriptors and how they impact alterations in the model’s response.

Similar to previously discussed methods, the interpretive complexity of these ensemble methods, or the identification of key descriptors, escalates as the dataset’s dimensionality increases. The types of descriptors and preprocessing strategies applied further influence this complexity, particularly when generating rules or identifying highly relevant features.

In contrast, probability-based methods introduce uncertainty into the prediction system. This facilitates the recognition of examples with a high probability of belonging to the predicted class or response, distinguishing them from other examples with lower probabilities. This augmentation enhances interpretability. Nevertheless, various statistical strategies and pattern identification techniques must be applied to comprehend probability’s effect fully. In synergy with feature analysis strategies, this approach enables the recognition of attributes playing a fundamental role in classifying examples and the identification of edge cases.

### 3.6 Incorporating XAI into protein properties predictions

AI systems, utilizing DL and ML algorithms, are crucial in addressing significant challenges in protein engineering Kouba et al. (2023). However, the complexity of DL or ML models often renders them opaque and perceived as *“black box”* systems, creating challenges in understanding the rationale behind their decisions (Samek et al., 2019). This lack of transparency poses difficulties for end-users, decision-makers, and ML developers (Arrieta et al., 2020). In sensitive domains like the design of therapeutic proteins and enzymes Kamerzell and Middaugh (2021) or therapeutic peptides Basith et al. (2020), the necessity for explainability and accountability is not only desirable but also legally mandated for AI systems that have substantial influence on human lives (Holzinger et al., 2022). Concerns regarding fairness have also arisen, emphasizing the need to prevent bias or discrimination in algorithmic decisions based on sensitive attributes.

Most state-of-the-art interpretable ML methods are domain-agnostic and have evolved from computer vision, automated reasoning, or statistics. This evolution makes their direct application to protein engineering and design problems challenging without customization and domain-specific adaptation (Kouba et al., 2023).

A perspective on XAI in the context of protein function prediction is provided by Le and Quoc (2023). The authors emphasized the importance of transparency in understanding predictions, particularly in the complex domain of protein functions. Cai et al. (2022) introduced an interpretable ML algorithm to predict disordered protein phase separation. The algorithm relied on biophysical interactions as critical features for its predictions. The goal was to enhance the understanding of the factors influencing protein phase separation, a phenomenon crucial to various cellular processes. The interpretability of the algorithm was emphasized, aiming to provide insights into the biological mechanisms underlying disordered protein phase separation. Bæk and Kepp (2022) discussed the estimation of protein mutant stability, explicitly focusing on the dataset and fitting dependencies involved in the modeling process. The primary objective was to develop simple, well-balanced, and easy-to-interpret models. The emphasis was on creating models that are not only accurate but also straightforward, ensuring a more accessible and comprehensible approach to estimating protein mutant stability (Bæk and Kepp, 2022). Bittrich et al. (2019) explored applying an interpretable classification model for examining early folding residues in protein folding. Bittrich et al. (2019) introduced an innovative adaptation of the Generalized Matrix Learning Vector Quantization (GMLVQ) machine learning technique. GMLVQ, a prototype-based supervised machine learning approach, offers extensive visualization capabilities and furnishes an interpretable classification model (Frank et al., 2016).

Hsu et al. (2022) developed an augmented model using evolutionary and assay-labeled data for protein fitness prediction. This model can be considered a surrogate model—it is ridge regression and sequence density evaluation encoded site-specific amino acid features. A Bayesian interpretation of this augmentation was provided, updating the prior from evolutionary and assay-labeled data, focusing on site-specific parameters. Furthermore, only one linear regression model was trained on a probability density model already in place.

## 4 Developing explainable predictive models for data-driven protein engineering: Pipelines and possible applications

Data-driven protein engineering is dedicated to devising and implementing computational methods to assist directed evolution and rational design strategies to enhance the properties of proteins or introduce new characteristics or functionalities of interest (Wittmund et al., 2022). Data-driven protein engineering primarily focuses on developing predictive models using AI techniques or statistical approaches (Siedhoff et al., 2021). The conventional pipeline for training predictive models in protein engineering, as illustrated in Figure **1**, encompasses stages addressing challenges in numerical representation, the training of generalizable predictive models, and model validation. ML methods leverage pre-existing mutational data to predict protein functions without relying on a detailed physical model. These approaches use available experimental and simulation data to aid in identifying and annotating promising enzyme variants. Furthermore, they contribute to suggesting beneficial mutations to improve known targets. Apart from classical directed evolution and (semi)rational design approaches, ML methods have been increasingly applied to find patterns in data that help predict protein properties such as enzyme stability, solubility, function, and substrate specificity and guide rational protein design strategies Kouba et al. (2023).

Despite this progress, the current predictive models’ transparency, interpretability, and explainability often go overlooked, leaving a gap in understanding how models learn or why they generate specific predictions. Bridging this gap can provide valuable insights to guide rational design methods or comprehend evolutionary design approaches (Kouba et al., 2023).

In the following section, we present and scrutinize various strategies and development pipelines for creating predictive models tailored for common protein engineering tasks, incorporating XAI approaches. We propose three predictive modeling pipelines that combine diverse protein representation techniques, predictive model training methods, and XAI strategies elucidated in the preceding sections of this work. Within each pipeline, we delve into distinct challenges to aid protein design, explaining potential avenues for acquiring knowledge that facilitates interpreting and clarifying the generated models. We aim to leverage this knowledge in guiding the design of proteins endowed with desirable properties. Subsequently, we design and implement two case studies: the classification of DNA-binding proteins and the prediction of green fluorescent proteins’ brightness. These case studies illustrate how integrating XAI methods can streamline the identification of rules on the target function or assist in designing proteins possessing sought-after properties.

### 4.1 Applying feature-attribution methods to explain *“black box”* predictive models

XAI methods, focused on characteristic analysis, offer a means to discern pertinent attributes based on the target response. The application pipeline for these strategies is illustrated in Figure **3** A. Initially, a protein characterization strategy is employed using descriptors, which may be derived from the protein sequence and structure. Additionally, descriptors encompassing physicochemical properties, thermodynamics, and phylogenetic and functional domain information are considered. Following a feature engineering process, the most informative descriptors concerning the problem are selected, and a predictive model is trained. Various strategies, including deep learning architectures such as LSTM or CNN-1D, are explored, often resulting in developing a high-performing yet complex *“black box”* predictive model.

**Figure 3:**
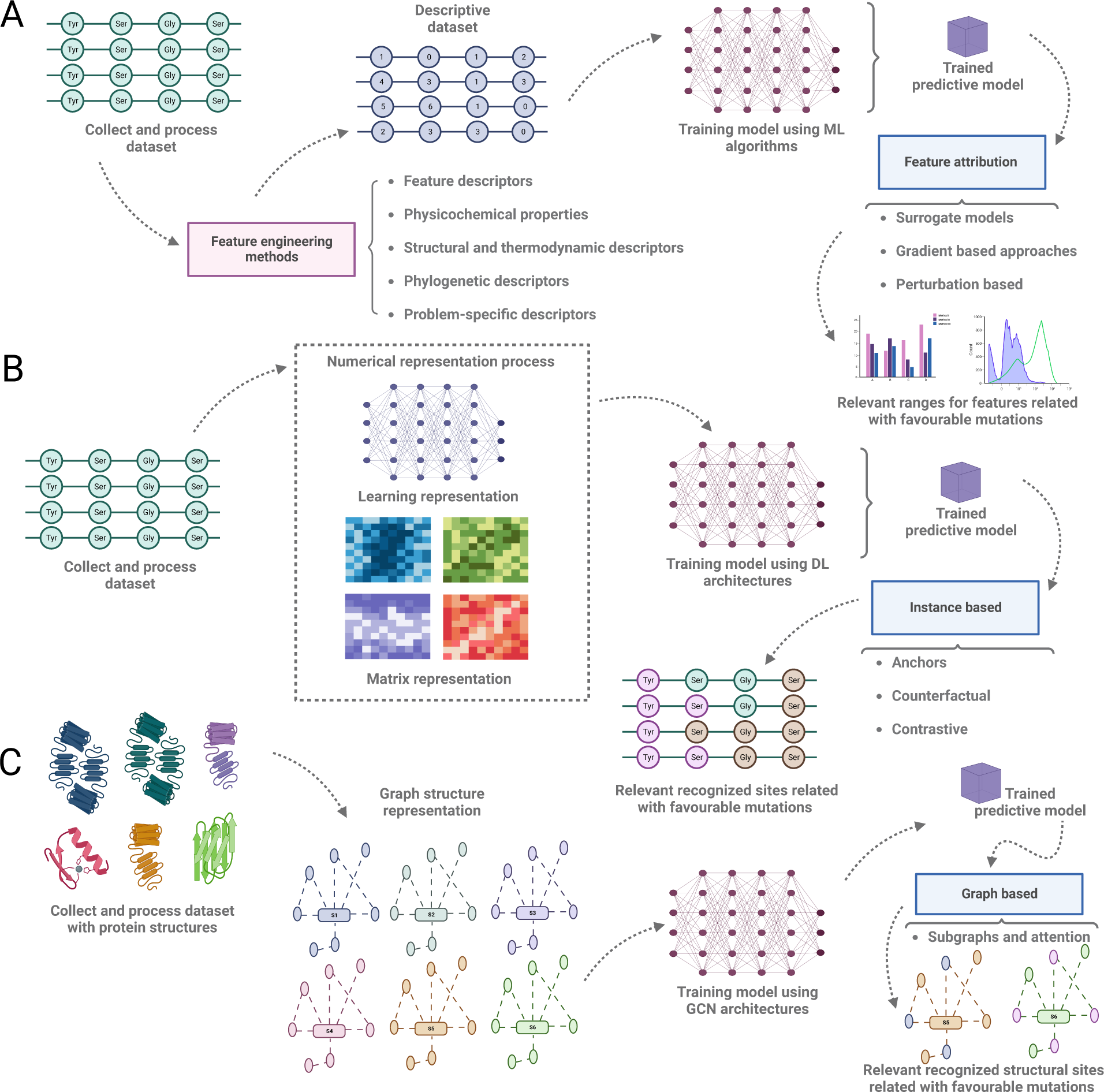
Proposed pipeline to develop explainable predictive model support by XAI strategies. **A** Pipeline to develop predictive models for protein engineering task support by a feature engineering approach to characterize protein sequences and feature attribution methods as XAI strategy, surrogate model facilitates the explainability of the *“black box”* model and gradient-based and perturbation-based support the identification of relevant features. **B** Relevant zones or site identification in protein sequences through instance-based approaches. The protein sequences are represented by learning representations or matrix strategies, and then DL architectures are employed to train predictive models. Finally, instance-based approaches detect relevant sites in the protein sequences as targets for site-directed mutagenesis. **C** Relevant structural zone identification by graph-based approach. First, the protein structures are represented as graph structures, and then a GCN architecture is trained to build the predictive model. Finally, subgraphs and attention methods are employed to detect relevant zones correlated with favorable properties as targets for rational design.

Simplification systems can be implemented using surrogate model techniques once the model is generated. This approach chooses simplified and interpretable ML algorithms like linear methods, decision tree strategies, or rule-based systems. The surrogate model is developed by randomly subsampling the original dataset, utilizing the trained predictive model to generate predictions, and using these predictions to label the subsampled examples. The resultant dataset is then employed to train the surrogate model. Interpretation of the surrogate model involves coefficient analysis (for regression methods), prediction differentiation via attributes, or evaluation of rules generated for the predictive system (for decision trees and rule-based models). This interpretation enables the extraction of insights into the predictive model, identifying rules or attribute values influencing tendencies to increase or decrease the property of interest associated with the predictions (Jiménez-Luna et al., 2020).

Alternatively, in developing predictive models based on deep learning architectures, methods utilizing gradients allow the identification of relevant attributes through weight mapping, considering the values of the gradients. Positive gradient values indicate higher relevance, while negative values suggest non-relevance for the prediction.

Finally, employing perturbation-based techniques allows the identification of the most sensitive attributes based on the model and its predictions. This facilitates the association of sensitivity with ranges, indicating where significant prediction changes occur for the property of interest. In conjunction with gradient-based methods, this approach aids in recognizing groups of attributes with high relevance and the value ranges where the model exhibits greater or lesser sensitivity.

Identifying the most relevant attributes through XAI techniques, particularly those focusing on feature attribution, streamlines the classification of attributes crucial to the predictions made by the model. This guides the rational design of proteins, directing necessary modifications to develop mutational variants that acquire the most relevant descriptive characteristics. It also helps restrict the ranges these descriptors can maintain or alter the property of interest. This approach benefits the design orientation by recognizing specific values for new mutational variants, effectively narrowing the search space.

### 4.2 Employing instance-based approaches to detect relevant sites in protein sequences

Instance-based methods play a crucial role in pinpointing key regions or sites within the representation space of datasets. When predicting a property of interest for a set of proteins, the typical workflow involves constructing predictive models using numerical representation strategies, such as representation learning, matrix representation methods, or color images. This process encompasses developing embeddings (the representation method), designing and implementing deep learning architectures, training predictive models, and evaluating model performance and validation. However, the resulting model often becomes a *“black box”*, lacking transparency and interpretability due to the representation via embedding and training through deep learning architectures (See Figure **3** B for more details).

Gradient-based methods can be employed to enhance model transparency and interpretability. Yet, achieving a clear causal relationship between the properties or characteristics of interest and the model’s predictions may be challenging. In such cases, instance-based methods come into play to identify relevant areas or sites in the representation. Among these methods, applying anchors is one approach for pinpointing specific sites contributing to the desired property or prediction values. These sites can be categorized as enhancing and diminishing the property. Alternatively, methods like counterfactuals enable stability identification by applying permutations. From a practical standpoint, this allows the analysis and reconstruction of evolutionary trajectories, indicating mutational changes that enhance the property of interest or introduce new characteristics.

Incorporating contrastive approaches, anchors, and counterfactuals further refines the identification of mutational targets by offering individual explanations and associating a sensitivity evaluation.

Across all instance-based strategies, the focus is on identifying relevant sites or areas in the data space crucial for prediction. Recognizing these areas allows classification or categorization based on their impact on improving or diminishing the property of interest. The inverse process of numerical representation can be complex, especially in learning representations or matrix methods without decoding techniques. However, if decoding techniques are available, mapping these areas and comparing them with the response of interest helps identify areas with a higher potential for improvement and those sensitive to changes.

This information can be cross-referenced with structural studies or evolutionary coupling to analyze feasibility from thermodynamic and phylogenetic perspectives. Alternatively, deep generative model methods can be incorporated or coupled, guiding sequence generation by marking specific detected areas of interest as positive and avoiding regions that harm the property.

### 4.3 Combining graph structure representations with graph-based approaches to detecting non-obvious structural sites

When utilizing graph structures to depict proteins, nodes are employed to represent atoms, residues, or centroids. At the same time, edges symbolize potential interactions assessed through distance methods or strategies involving weak electrostatic interactions. Alternatively, various descriptors can characterize the nodes, whether atomic or residue descriptors, contingent on the specific representation. Once proteins are transformed into graphs, implementing architectures based on Graph Convolutional Networks (GCN) or similar becomes imperative for training predictive models. Figure **3** C outlines a pipeline for crafting predictive models for protein engineering tasks using GCN methods.

However, developing predictive models based on GCN inherently yields a *“black box”* model that defies straightforward interpretation. Graph-based methods come into play to unravel and expound upon the model. Specifically, subgraph-based methods indirectly facilitate the identification of structural zones (hot spots) in proteins that significantly influence predictions. This identification allows for classification based on the impact on the property, pinpointing whether it resides in a location pertinent to the protein’s function, such as an active or allosteric site in the case of enzyme engineering. This recognition of non-obvious structural areas empowers the orientation and guidance of structure-based variant design, encompassing modifications to pockets, insertion or removal of residues, and evaluation of more intricate alterations from a rational design perspective.

However, a primary challenge encountered when employing a GCN-based pipeline is the necessity for the protein’s structure every time new examples are explored. This requirement is limited, especially in massive explorations, due to the associated computational costs, the selection of protein structure prediction strategies, and the evaluation and validation methods employed for the predicted structures.

### 4.4 Self-training and ensemble learning to interpret and explain complex predictive models

Self-explaining methods aim to produce interpretable predictions by aligning with the design and structure of the models. In contrast, uncertainty estimation methods focus on assessing probabilities to manage error margins around predictions. Combining both approaches into a unified predictive/explanatory system addresses the challenges of constructing high-performance predictive systems with efficient explanations. The resulting explanations can guide protein design by providing insights into the rules governing distinctions between predictions from the model.

Following a conventional numerical representation strategy, amino acid coding methods utilize physicochemical properties based on the descriptors proposed in (Medina-Ortiz et al., 2022), generating eight sets of coded datasets. Each coded dataset is trained with an exploratory approach, evaluating traditional ML methods and DL architectures. The optimal methods for each physicochemical property are selected using statistical techniques, and an ensemble strategy is devised to maximize system performance by combining predictive models and coding strategies.

Combining self-explaining and uncertainty estimation methods is recommended as an XAI strategy. For self-explanation, prototype-based approaches or concept learning can be employed to identify abstract elements (prototypes or concepts, depending on the chosen strategy) that are descriptive and representative based on the response of interest. These abstract elements serve as representative subsets guiding the design of proteins with tunable properties, enabling a focus on common properties and characteristics shared among them. Alternatively, applying natural language explanations helps identify descriptive rules for interpreting predictions based on specific points or areas of interest in the encoded protein vector. Regardless of the chosen approach, decoding systems are required to interpret potential changes from the sequence.

Coupling self-explaining methods with uncertainty estimation makes it possible to develop estimators associated with predictions. Typically based on the use of Bayesian distributions, these probabilities allow the filtering of predictions with a high likelihood of success from those with percentages with low probability. Filtering examples with a high probability of success, representing favorable changes in the property of interest based on the native protein, can be employed as a support system for validating and exploring new variants. These variants are guided by relevant areas or sites identified in the previous step. This approach, developing a predictive/explanatory system with hybrid techniques, encourages support and design guidance for mutational variants with desirable properties and a high success rate.

### 4.5 Demonstrating how to incorporate XAI approaches in predictive models

Two predictive models were meticulously crafted to exemplify the practical integration of XAI strategies in data-driven predictive models. The initial model delineates a functional protein classification system with a DNA-binding function. The second model delves into a predictive methodology for estimating average brightness in the green fluorescent protein (GFP) (See Section S1 in Supplementary Materials for more details). In both instances, the employed predictive model training strategies combine amino acid encoding techniques leveraging physicochemical properties, the exploration of machine learning algorithms rooted in supervised learning, and the reasonable selection of optimal algorithms through performance evaluation metrics (see Section S2 in Supplementary Materials for more details). After rigorous training and evaluation of the predictive models, XAI techniques were employed to decipher and elucidate the model-generated predictions (See Section S3 in Supplementary Materials for more details). Simultaneously, alternatives were assessed to assimilate the acquired knowledge in bolstering the design of proteins endowed with desirable properties, contingent on the tasks considered in predictive evaluations. The results for the DNA-binding classification system and the average brightness in the protein landscape for GFP protein are described in the following sections.

#### 4.5.1 Interpreting and explaining the DNA-binding classification system

To construct DNA-binding classification systems, nineteen supervised learning algorithms were investigated, utilizing the *α*-structure property as input to encode protein sequences (refer to Section S2 of Supplementary Materials for detailed information). The XGBoost algorithm demonstrated superior performance, achieving a precision of 77% and recall of 0.77. Notably, the chosen model operates as a *“black box”* model, necessitating applying XAI strategies for interpretation.

To do this, surrogate model-based analyses were initially employed, utilizing decision tree-based algorithms to derive simplifications. The replicability of the surrogate model was outstanding, with metrics nearing 65% precision and accuracy. However, a granular analysis of decision rules revealed that positions 1, 4, 14, 12, 41, and 7 played pivotal roles in division or separation functions. Despite the high replicability, the interpretation of specific functions and support for changes or mutational variants contributing to DNA-binding function proved challenging due to the high dimensionality of the applied encoders.

Alternatively, feature-based methods such as SHAP and LIME were employed to identify relevant positions based on the activity of interest. Figures **4** A and **4** C present SHAP analyses for the 4BLF and A4SLN8 proteins, both classified as DNA-binding. Notably, physicochemical property values at positions 151, 129, and 296 exhibited high relevance in attributing DNA-binding function to unknown proteins. Positions 2, 267, and 135, on the other hand, strongly favored the classification as Non-DNA-binding. Consistent results were obtained through surrogate model methods and LIME analyses (refer to Section 4 in Supplementary Materials).

**Figure 4:**
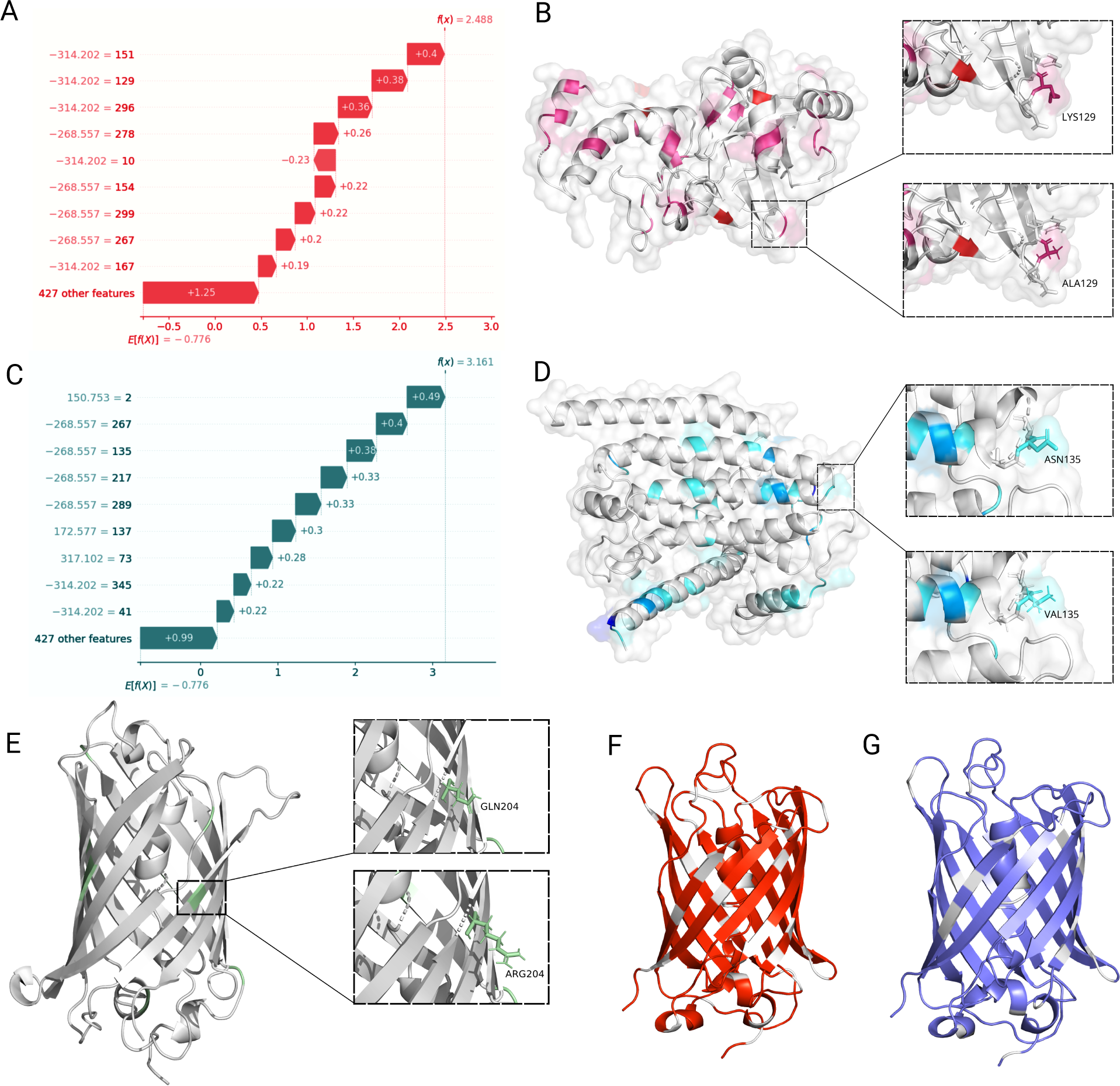
Incorporating XAI strategies for classification and predictive models to facilitate interpretability, explainability, and assist the protein design. **A** SHAP analysis for the 4BLF protein sequence classified as a DNA-binding protein. **B** Structural sites of the 4BLF protein with a higher tendency to cause a loss of function when mutated individually or in combination. **C** SHAP analysis for the A4SLN8 protein initially classified as non-DNA-binding. **D** Structural sites of the A4SLN8 protein that, when mutated, causes a functional change, acquiring the ability to bind to DNA. **E** Structural sites detected through counterfactual methods to improve the median brightness values. **F** Mutational variant with sites that cause a loss of function and a decrease in average brightness compared to the native protein. **G** Mutational changes to GFP involve an increase in median brightness but a loss of function.

SHAP analyses, coupled with counterfactual strategies, not only facilitated the examination of model interpretability by pinpointing changes affecting the binding or recognition site but also illustrated the structural zones in the 4BLF protein where mutations commonly result in an unfavorable change or loss of function (Figure **4** B). For instance, a specific change at position 129 (lysine to alanine) directly caused a loss of function in the DNA-binding protein.

Furthermore, these strategies supported the identification of rules enabling the acquisition of DNA-binding function, laying the groundwork for designing proteins with desirable properties. Figures **4** B and **4** D detail substitutions and modification zones that influence function, providing valuable insights for protein design.

While these methodologies offer significant advantages regarding explainability and interpretability, their application is contingent on representation strategies. Although effective in classifying proteins with DNA-binding functions, using physicochemical properties for numerical representation reveals limitations in exploring latent spaces. This constraint hinders the insertion and elimination of residues, highlighting a considerable opportunity for improvement in assisting the design of proteins with desirable functions.

#### 4.5.2 Guiding protein design supported by XAI approaches exemplified with GFP proteins

To illustrate the significant impact of ML methods on designing proteins with desirable properties, we showcase how protein modifications can either enhance or impair a specific property. Concurrently, we demonstrate the feasibility of preserving or improving the property at the expense of loss of function, emphasizing the intricate nature of designing mutational variants with desirable properties. This underscores the imperative need for a collaborative approach involving predictive systems, functional classification methods, thermodynamic studies, and the application of bioinformatics tools to validate mutational variants proposed by predictive systems.

To do this, a regression model was initially developed to estimate the average brightness in the mutational landscape of GFP Sarkisyan et al. (2016). The presented model achieves a Pearson coefficient of 0.7 and an MSE of 0.4 as performance metrics (See section S2 in the Supplementary Materials for more details). Recognized as a *“black box”* model, it is an ideal input for explainability analyses. We employed feature-based XAI strategies to study the average brightness value by identifying key amino acid positions and residues. SHAP analysis and individual queries in LIME revealed the relevance of positions such as 24, 107, 204, and 97 in improving the average brightness value (See Section 4 of Supplementary Materials for more details). However, it is crucial not to assume that modifications to these residues alone will invariably enhance a property, as various factors necessitate analysis before validating proposed substitutions. Notably, these positions are proximate to or associated with the functional domain of the proteins, suggesting that alterations in these residues impact both the property and the function. Furthermore, SHAP values and points of interest from LIME analyses elucidate these phenomena using information extracted solely from the model, generating potential ranges for property improvement. These ranges, representing values of physicochemical properties, can guide changes based on the employed encoder. However, this simplicity is attainable and interpretable primarily using amino acid encoding strategies as a numerical representation method.

After applying feature-based approaches, counterfactual strategies were employed to evaluate how perturbations could positively or negatively affect the average brightness value. In all cases analyzed, the average brightness value for the query variant serves as the baseline, and changes within a window of ±3 are explored.

First, positive effects were identified and analyzed using the counterfactual approaches. Figure **4** E shows mutations in the native protein that positively affect the median brightness values supported by the predictive models. In this case, nine mutations were introduced to improve the median brightness average, including V12N, E17G, and G145Y. As a result of the applied mutations, the median brightness values increase from 1.3 to 2.58. Most changes that positively affect the median brightness values are related to modifications of hydrophobic residues for hydrophilic or charged amino acids.

Alternatively, the same approach was applied to the variants with the lowest and the highest median brightness. This analysis aims to illustrate how unguided alterations in the protein can modify the median brightness value with a function loss. Figure **4** F shows a mutational variant with adverse effects on median brightness obtained from counterfactual analyses and a loss of function (identified via BLAST alignment). Conversely, lastly, Figure **4** G showcases a variant from the counterfactual analysis that improves the median brightness value but results in a loss of function.

Identifying mutational variants via counterfactual techniques, influencing the property of interest positively or negatively, underscores the considerable potential of XAI techniques as support for experimental protein design. However, additional considerations are necessary, involving strategies for forbidden or harmful functional zones, including checkpoints such as function evaluation, thermodynamic analyses, and potential stereochemical effects negatively impacting protein stability. This necessitates integrated computational systems promoting active learning of machine learning systems, incorporating experimental knowledge to devise an autonomous learning system guided by transfer learning or similar approaches.

## 5 Remaining Challenges and Future Opportunities

Designing proteins with desired properties has been a significant challenge in protein engineering with implications in biotechnology and bioengineering. AI advances have greatly enhanced data-driven protein engineering techniques. Fusing predictive ML models with experimental approaches facilitates the reconstruction of mutational landscapes and the navigation of latent spaces.

Different strategies can be applied depending on the type of response and the information used to train predictive models. These strategies combine methods for characterizing or numerically representing protein sequences with machine learning algorithms or deep learning architectures. Most strategies for training predictive models use numeric representation methods focused on characterization or encoding strategies, along with classic supervised learning algorithms such as Random Forest, SVM, or Decision Tree.

Alternatively, representation methods based on matrix information or graph structures for proteins have been applied in more complex predictive algorithms. Complex DL architectures based on convolution or graph methods are required when employing these representation strategies. This increases the complexity of the models and raises non-linearity, high dimensionality, and low interpretability and explainability of the models in exchange for improved predictive performance and greater generalization.

Recent methods focus on using large language models, defined as protein language models, promoting transfer learning. The embedding generated by pre-trained models of protein language models is also utilized with deep learning architectures such as CNN, LSTM, and hybrids. Although these methods significantly enhance performance compared to traditional approaches, facilitating the efficient reconstruction of mutational landscapes, they increase complexity regarding model transparency and interpretability. Furthermore, if methods facilitating navigation of latent spaces are desired, architectures enabling this must be introduced, involving autoencoders, variational autoencoders, or deep generative model methods.

In this context, advances in artificial intelligence methods applied to protein engineering have focused on improving predictive model performance, facilitating efficient reconstruction of mutational landscapes, and promoting navigation in latent spaces to guide protein design. However, this has increased model complexity, resulting in a lack of interpretation and explainability, hindering understanding of how the model operates to address questions such as why and how inspiring the application of XAI methods.

XAI methods have addressed this gap by focusing on developing computational methods and strategies to comprehend and elucidate predictions generated by these models. Techniques such as feature attribution, instance-based analysis, graph-based methodologies, self-explaining models, and uncertainty estimation have been developed, each catering to distinct perspectives. The choice of method depends on the descriptors or representation strategies and the predictive modeling methods employed. Regardless of the strategy, comprehending how predictive models learn facilitates a deeper understanding of the underlying problems, enabling the resolution of design and evaluation challenges in diverse fields such as biotechnology, medicine, and pharmacology.

In protein design and engineering, XAI methods are crucial in understanding the necessary changes to introduce new functionalities or properties. This knowledge can guide protein design by identifying the key characteristics describing functional properties, aiding rational design approaches. Understanding the mechanisms that prompt a change in category or class can indirectly shed light on evolutionary processes, benefiting techniques like directed evolution. This synergy between experimentalists and data scientists opens avenues for ML-assisted protein design.

Nevertheless, applying XAI techniques to protein engineering poses challenges concerning representation strategies. Current leading strategies, such as protein language models, require explanatory models and interpretations of non-linear relationships within sequence embeddings. Physicochemical property coding may offer a more straightforward interpretation, but decoding methods are essential. Graph-based techniques, leveraging graph convolutional network (GCN) architectures, can address complex problems like protein-protein interactions, but their computational costs pose challenges. Although subgraph identification or attention-based techniques in graph-based XAI methods are feasible, understanding and characterizing identified areas remain crucial.

Interdisciplinary collaboration is encouraged to bridge the gap between mathematical representations and algorithms, and domain-specific knowledge for XAI models has unveiled challenges and promising avenues for future research. These challenges include adapting to complex models, integrating symbolic and numerical representations, providing dynamic explanations, ensuring scalability, and establishing benchmarks. Some work will need to be done on adaptable algorithms, holistic integration methods, scalability solutions, user-centric evaluations, and interdisciplinary collaboration. In the meantime, high-dimensional data representations, improved interpretability, tailored algorithms, new evaluation metrics, and visualization techniques are highly recommended to be considered in future works.

Developing efficient algorithms and representations to handle the growing size of datasets and models is the main part of the scalability solution that can enable real-time explanations without computational bottlenecks.

Dealing with diverse and complex information requires incorporating domain knowledge into mathematical representations. Establishing standardized benchmarks is equally imperative to evaluate the effectiveness of different mathematical representations across diverse XAI approaches on a common basis. Meanwhile, the innovative and holistic methods leveraging the seamless integration of symbolic and numerical representations enhance interpretability and transparency in complex AI architectures.

Designing algorithms that can scale and dynamically adapt to complex models’ intricacies can be an interesting topic for future work. The resulting algorithms will address the dynamic nature of evolving data distributions and model behaviors. Considering the seamless blending of symbolic and numerical representations to provide comprehensive and informative explanations, holistic integration methods take center stage. Additionally, user-centric evaluations enable end-users to interpret and trust XAI explanations.

Comprehensively assessing and communicating the performance and interpretability of AI models can be achieved by striving for new evaluation metrics and visualization techniques. Creating new representations specifically tailored for high-dimensional data, improving the interpretability of existing algorithms, and developing algorithms tailored to specific applications are other topics for future work.

In conclusion, integrating XAI methods into predictive modeling for protein engineering faces several challenges. Yet, comprehending the learning process and identifying decision rules of predictive models enriches knowledge, offering substantial support for protein design. This tool can potentially address biotechnological challenges and uncover abstract evolutionary rules.

## Author contributions statement

All authors have read and agreed to the published version of the manuscript.

## Codes and data availability statement

Established learning and validation sets of applied data sets are given in Supporting Information. The source code to replicate all exploring methods and XAI strategies, including Jupyter Notebook to facilitate the execution of the different techniques, is publicly available on GitHub (https://github.com/ Protein-Engineering-Framework/XAI_Protein_Engineering) under MIT License. All source code was written in Python 3 and developed to be run using the Jupyter Notebook examples.

## Acknowledgments

DM-O acknowledges ANID for the project “SUBVENCIÓN A INSTALACIÓN EN LA ACADEMIA CONVOCATORIA AÑO 2022”, Folio 85220004. DM-O gratefully acknowledges support from the Centre for Biotechnology and Bioengineering - CeBiB (PIA project FB0001, Conicyt, Chile). DM-O gratefully acknowledges Nicole Soto Garćıa for her support during the image development. MDD acknowledges funding by the Deutsche Forschungsgemeinschaft (DFG, German Research Foundation) - within the Priority Program Molecular Machine Learning SPP2363 (Project Number 497207454). MDD acknowledges EU COST Action 841 CA21162 (COZYME) for support.

