## Supplementary Information for "Interpretable and explainable predictive machine learning models for data-driven protein engineering"

[1]Departamento de Ingeniería En Computación, Universidad de Magallanes, Avenida Bulnes 01855, Punta Arenas, Chile.
[2]Centre for Biotechnology and Bioengineering, CeBiB, Beauchef 851, Santiago, Chile
[3]Department of Bioorganic Chemistry, Leibniz Institute of Plant Biochemistry, Weinberg 3, 06120 Halle, Germany

**S4 Explainable artificial intelligence results** **6**

*

### S1 Summary of dataset and tasks employed to train predictive models

For the presented work, two datasets were employed to train predictive models and to explore the different Explainable Artificial Intelligence (XAI) approaches. Table **S1** describes the datasets in terms of the number of examples, tasks, and sources.

**Supplementary Table S1**: Description of the dataset employed to train and explore XAI approaches

| # | Task | Description | Examples | Sources |
|---|---|---|---|---|
| 1 | DNA-Binding protein identification | Dataset with protein sequences classified as a DNA-Binding protein(10070 examples) and non-DNA-Binding protein (13739 examples). | 23809 | (Mishra et al., 2019) |
| 2 | Estimation of brightness in GFP protein | Local fitness landscape of the green fluorescent protein. A dataset with different variants sequences with a wide range of number of mutations, including one point mutation until 15 mutations at the same time. | 51714 | Sarkisyan et al. (2016) |

In the case of DNA-Binding protein classification, the objective was to build a binary classification model. In contrast, the aim is to train a regression model to estimate brightness in GFP protein (green fluorescent protein).

### S2 Strategies to train predictive models through machine learning

A classical data-driven pipeline was used to train predictive models for the DNA-binding classification system and estimate brightness in the GFP landscape. First, a dataset preprocessing was performed in the case of the DNA-binding classification system. These preprocesses implied a filter strategy to remove unlabeled examples and protein sequences with extreme length values. The length filter was conducted to facilitate the interpretation and explanation during the decoding processing. Also, to prevent noise during the *zero-padding* process, ensure encoded vectors are the same length.

Once the preprocessing stage is finished, the amino acid encoding strategy is applied. For both datasets, a physicochemical encoding method employing the $\alpha-$structure encoder property as proposed in (Medina-Ortiz et al., 2022). The selection of the specific physicochemical properties was random, and the amino acid encoding approach was based on facilitating the decoder process. In the case of the DNA-binding dataset, a $zero-padding$ method was employed to standardize the size of the generated vector during the numerical representation process.

Then, a classical pipeline to train predictive models supported by a machine learning approach was applied for both evaluated tasks in the presented work (Medina-Ortiz et al., 2020b; Siedhoff et al., 2020). First, the dataset is split randomly into training and validation datasets in a 70:30 proportion. Then, using the training dataset, different supervised learning algorithms are applied, including Support Vector Machine (SVM), Random Forest, Bagging, Adaboost, and various *"black box"* models. During the training process, a $k-$fold cross-validation ($k = 10$) was applied to prevent the overfitting. To finish, the validation dataset was employed to obtain the performances of the trained predictive models. The classification models utilized accuracy, precision, recall, and f-score metrics. In contrast, the regression models used metrics like the Pearson correlation coefficient and Root Means Squared Error (RMSE).

Once the performances are obtained, a simple selection process is generated to choose the models for applying the XAI approaches. First, the machine learning algorithm needs to be classified as a black-box model, and the performances for the selection models need to be high.

Seventeen supervised classification algorithms were explored for the DNA-binding classification model. Table **S2** describes the different validation performances. The best model based on all metrics was obtained by applying the XGBoost algorithm, achieving 77% of precision and a recall of 0.77. This model was selected to apply XAI strategies.

The XGBoost classifier for DNA-binding identification is described in Figure **S1**. The confusion matrix shows a high True positive rate for correctly identifying DNA-binding proteins. In contrast, the model achieves

**Supplementary Table S2**: Performances for DNA-binding classification models

| Description | Accuracy | F1 score | Precision | Recall |
|---|---|---|---|---|
| XGBoost | **0.772505** | **0.769346** | **0.771854** | **0.772505** |
| NuSVC | 0.757245 | 0.752464 | 0.757214 | 0.757245 |
| RandomForestClassifier | 0.755425 | 0.739444 | 0.777716 | 0.755425 |
| ExtraTreesClassifier | 0.745765 | 0.728046 | 0.768635 | 0.745765 |
| SVC | 0.742685 | 0.732074 | 0.749054 | 0.742685 |
| GradientBoostingClassifier | 0.722526 | 0.705372 | 0.735208 | 0.722526 |
| BaggingClassifier | 0.717626 | 0.706192 | 0.720433 | 0.717626 |
| KNeighborsClassifier | 0.692706 | 0.674708 | 0.697656 | 0.692706 |
| AdaBoostClassifier | 0.666387 | 0.659865 | 0.661413 | 0.666387 |
| DecisionTreeClassifier | 0.653507 | 0.654324 | 0.655455 | 0.653507 |
| LogisticRegression | 0.631947 | 0.626351 | 0.625906 | 0.631947 |
| LinearDiscriminantAnalysis | 0.630267 | 0.620834 | 0.622563 | 0.630267 |
| RidgeClassifier | 0.630127 | 0.620387 | 0.622330 | 0.630127 |
| GaussianProcessClassifier | 0.582528 | 0.429455 | 0.756909 | 0.582528 |
| SGDClassifier | 0.562929 | 0.563477 | 0.564093 | 0.562929 |
| QuadraticDiscriminantAnalysis | 0.557049 | 0.534742 | 0.646377 | 0.557049 |
| PassiveAggressiveClassifier | 0.550609 | 0.552707 | 0.556197 | 0.550609 |
| GaussianNB | 0.503010 | 0.465569 | 0.591879 | 0.503010 |

a 67% True negative rate, demonstrating problems in detecting non-DNA-binding problems. Alternatively, the ROC curve is calculated and processed during the training process. The AUC score, on average, is 0.85, showing excellent performances when increasing the false positive rate.

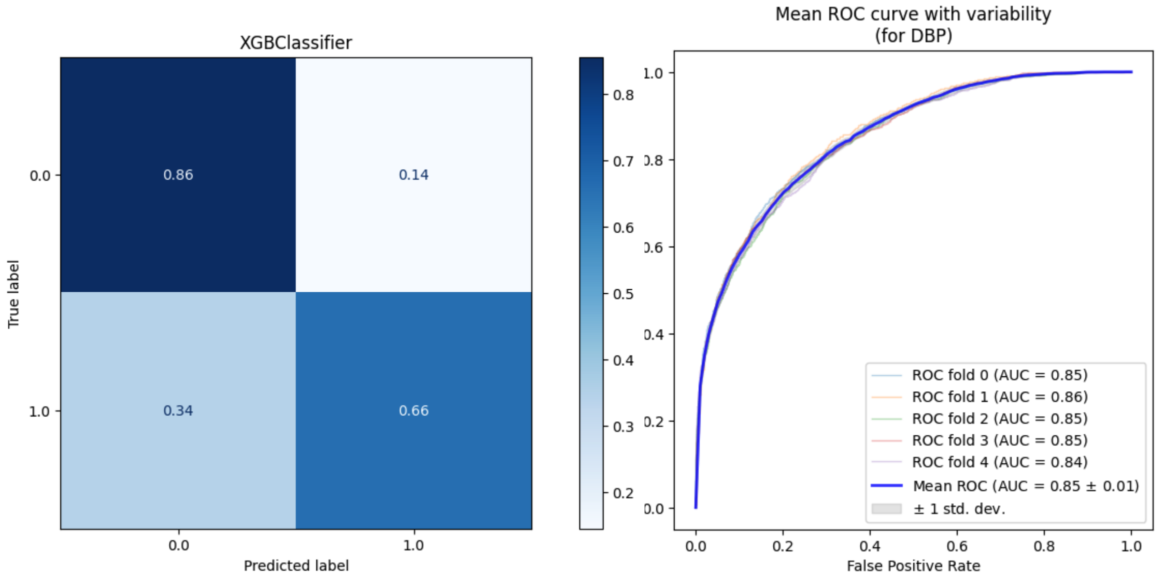

**Supplementary Figure S1**: **Descriptive plots for DNA-binding classification model**. (Left) Normalised confusion matrix for validated dataset (0 represents positive class and 1 represents negative class). The XGBoost model shows high True Positive values and some problems recognising harmful non-DNA-binding proteins with a True Negative value of 0.66. (Right) Mean ROC curve for DNA-binding classification model generated during the training process applying a $k-$fold cross-validation method to prevent over-fitting. The XGBoost classifier does not show significant differences during the cross-validation iterations.

In the case of the brightness predictive system's case, eleven predictive models were generated by applying supervised predictive algorithms. The best model was the HistGradientBoostingRegressor, achieving a performance of 0.7 Pearson's coefficient and an MSE of 0.57. This model was selected to apply XAI strategies.

**Supplementary Table S3**: Performances for brightness prediction in GFP proteins

| Description | Pearson correlation | Mean abs error | Mean squared error |
|---|---|---|---|
| HistGradientBoostingRegressor | **0.704007** | **0.448933** | **0.576353** |
| RandomForestRegressor | 0.672724 | 0.379971 | 0.606045 |
| BaggingRegressor | 0.661961 | 0.381965 | 0.615930 |
| ExtraTreesRegressor | 0.622625 | 0.391116 | 0.650780 |
| DecisionTreeRegressor | 0.505602 | 0.420902 | 0.744879 |
| GradientBoostingRegressor | 0.369736 | 0.778566 | 0.841025 |
| SVR | 0.303581 | 0.631269 | 0.884063 |
| KNeighborsRegressor | 0.103962 | 0.691290 | 1.002792 |
| AdaBoostRegressor | 0.026406 | 1.005113 | 1.045289 |
| PassiveAggressiveRegressor | -3.068658 | 1.960563 | 2.136846 |
| GaussianProcessRegressor | -5.723986 | 2.488863 | 2.747015 |

The HistGradientBoosting regressor model selected to apply and explore XAI strategies is described in Figure **S2**. First, the dataset is analysed using a boxplot (See Figure **S2** A) and a histogram (See Figure **S2** B). The median brightness shows a bi-modal behaviour. This distribution negatively affects the generalisation of the trained model. More than 25% of the examples have median brightness values near 1.3. In contrast, more than 50% of the examples have values over 3.5, and only a few examples have median brightness values between 1.3 and 3.5.

The trained model shows preferences to predict only high and low values and a low capability to predict examples with middle values (See Figure **S2** C). This behaviour is due to the input values used to train the predictive model. Despite achieving high performance related to Pearson's coefficient and low RMSE values, the model tends to overestimate values. It is possible to solve problems by using more complex strategies to represent the protein sequences numerically and more complicated methods to train predictive models that show problems to be solved. However, the trained model is only trained to apply XAI methods.

### S3 Machine learning and XAI implementation details

All source code implemented in this work was constructed using the Python v3.9.16 programming language. The machine learning predictive models were trained using the supervised learning algorithms in the DMAKit library (Medina-Ortiz et al., 2020a), for the numerical representation strategies was implemented an amino acid physicochemical encoding method employing the $\alpha-$structure encoder proposed in (Medina-Ortiz et al., 2022). Finally, the trained models were exported as joblib instances to facilitate usability.

In the case of XAI, strategies were explored through instance-based and feature-based approaches. In the case of the feature-based approach, the following strategies were implemented:

#### S3.1 Local Interpretable Model-agnostic Explanations

Local Interpretable Model-agnostic (LIME) is a technique used in machine learning for model interpretation and explanation. The main goal of LIME is to provide insights into the predictions of complex machine learning models, making them more interpretable and understandable, especially for individual instances or predictions (Jiménez-Luna et al., 2020).

In practical terms, LIME generates locally faithful explanations for individual predictions. It does so by perturbing the input features around a specific instance, obtaining predictions from the black-box model for these perturbed instances, and then fitting a simpler, interpretable model (such as linear regression) to approximate the behavior of the complex model in the local region around the instance in question (Ribeiro et al., 2016). LIME helps bridge the gap between the inherent complexity of advanced machine learning models and the need for human-understandable explanations, making it a valuable tool for model interpretability and explainability (Gevaert et al., 2022).

In this work, the lime Python package (Ribeiro et al., 2016) was employed to apply the Lime method in the case of classification systems. Different positive (DNA-binding protein) and negative (non-DNA-binding protein) query points were generated and evaluated to determine each point's top ten more relevant features. The same package and strategies were applied for regression tasks related to the brightness GFP landscape predictions.

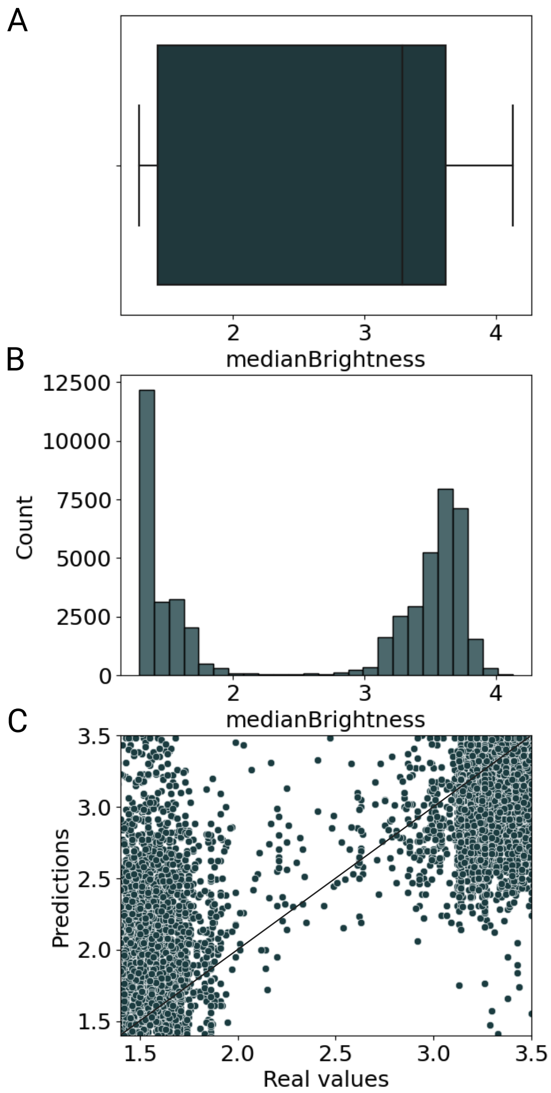

**Supplementary Figure S2**: **GFP dataset exploration and evaluating predictive models for median brightness predictions using HistGradientBoosting algorithm**. **A** Boxplot distribution for the median brightness values in GFP dataset. **B** Histogram for the median brightness values in GFP dataset. There are a clear bi-modal distribution in the dataset considering high and low median brightness values. **C** Scatter plot for HistGradientBoosting model plotting the real values v/s the predictions generated by the model.

#### S3.2 Surrogate model

A surrogate model, in the context of machine learning and optimization, is a simplified and interpretable model that approximates the behavior of a more complex or computationally expensive model. The purpose of a surrogate model is to serve as a stand-in for the original model, offering a quicker and less resource-intensive alternative, especially when the original model is time-consuming or expensive to evaluate (Elton, 2020). Using surrogate models is a strategy to balance the trade-off between model accuracy and computational efficiency, mainly when dealing with models that are challenging to evaluate directly (Elton, 2020).

A decision tree algorithm with a default hyperparameter was selected for classification and regression tasks. The training process was the same as explained previously, including the preparation dataset, hyperparameter selection, and validation step. Different strategies were implemented to evaluate the replicability of the surrogate model. In the case of the classification systems, classic metrics like accuracy, precision, and recall were employed. In contrast, R-squared metrics were used for the regression models.

#### S3.3 SHapley Additive exPlanations (SHAP) analysis

SHapley Additive exPlanations (SHAP) is a framework and a set of values used to explain the output of machine learning models. It is based on cooperative game theory, precisely the concept of Shapley values, which assigns a fair contribution to each player in a coalition game. SHAP analysis has become a popular tool for interpretable machine learning, enabling practitioners and researchers to understand the impact of each feature on individual predictions, thereby enhancing transparency and trust in machine learning models (Lundberg and Lee, 2017).

The SHAP analysis was implemented using the shap Python package (Lundberg and Lee, 2017). Different analyses were evaluated using the SHAP strategy. First, global analysis was explored to evaluate the most relevant values for physicochemical properties and their relations with the positions in the protein sequence. Then, Different queries were made to evaluate positive or negative functions and high, middle, and low brightness in the case of the GFP protein landscape.

In the case of the instance-based approach, the counterfactual method was explored. The dice_ml Python package implemented the counterfactual methods (Mothilal et al., 2020). The same strategy was generated for classification and regression tasks. In this strategy, ten counterfactuals were explored for positive and negative queries (in the case of the classification system) and a range of possible predictions generated by the predictive model (in the case of the regression system).

### S4 Explainable artificial intelligence results

Feature-based and Instance-Based approaches were employed to showcase the explicability and interpretability of the proposed models for identifying proteins with DNA-binding function and predicting brightness in the GFP protein landscape. Figure **S3** illustrates the outcomes of both global analyses and individual queries using the SHAP strategy, applied to classification models **S3** (A, C, E, G, I) and regression models **S3** (B, D, F, H, J).

In classification models, positions 1, 4, 14, 12, 41, and 7 consistently demonstrate the highest average SHAP values, signifying their greater relevance compared to other positions. Specifically, position 185 emerges as crucial for proteins manifesting the activity. In-depth analyses reveal that this residue corresponds to polar properties, distinguishing it from proteins lacking the function, which feature residues with hydrophobic characteristics. Notably, positions relevant to the negative query differ from those identified as pertinent in the positive query. This underscores an insightful interpretation of properties and residues concerning the function.

In the case of regression models, SHAP analyses reveal that positions such as 24, which corresponds to an asparagine residue in the wild-type protein, need to be maintained to preserve their function. This is interpreted in the global analysis and individual queries, showing high importance according to the SHAP analysis. Additionally, positions like 204 and 107 significantly impact the high and low brightness values prediction, respectively. This indicates a clear interpretation based on the original values of these residues, which correspond to threonine and tyrosine, respectively.

To enhance the SHAP analyses, we employed a LIME analysis strategy. Figure **S4** (A, B)illustrates the outcomes of queries generated for the DNA-binding classification model. In contrast, Figure **S4** (C, D) presents the results for queries associated with the brightness values estimation model. In classification models, queries

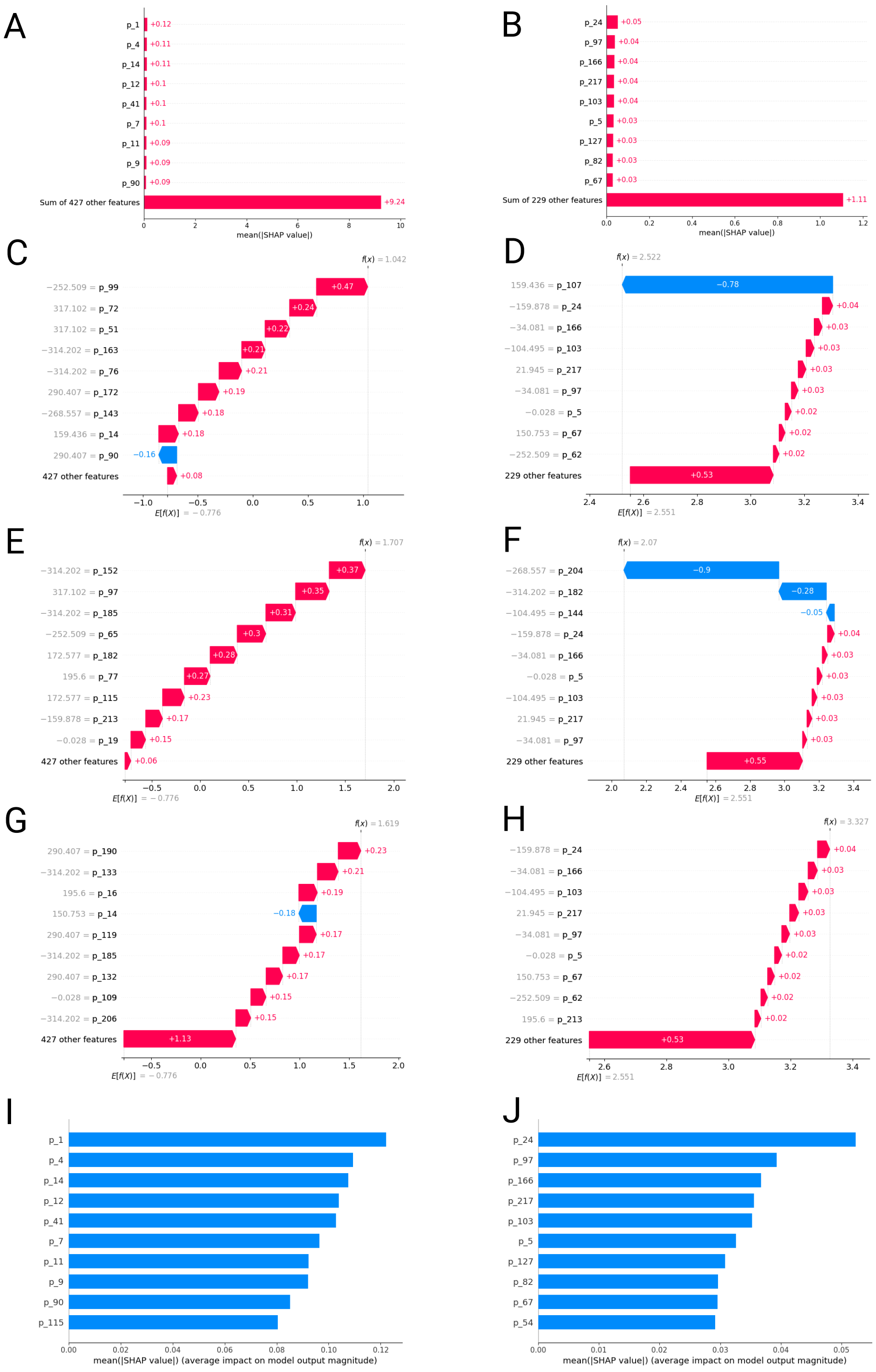

**Supplementary Figure S3**: **SHapley Additive exPlanations (SHAP) analysis for classification and regression models explored in this work**. **A** Global summary of the DNA-binding classification model's main positions and physicochemical properties. **B** Global summary of the main positions and physicochemical properties for GFP regression model. **C, E, G** Single instances of SHAP values for DNA-Binding classification model. **D, F, H** Single instances of SHAP values for GFP regression model. **I** The ten most relevant positions and physicochemical values for the DNA-binding classification model. **J** The ten most relevant positions and physicochemical values for the GFP regression model.

were conducted by evaluating positive and negative predictions. On the other hand, predictive models were queried based on high or low brightness values.

In the context of classification models, positions like 41 again assume a pivotal role, underscoring the importance of maintaining this position in terms of residues and properties to preserve the function. Meanwhile, positions 5, 75, and 76 maintain their significance for predictive models, corroborating the findings obtained through the SHAP analysis methods.

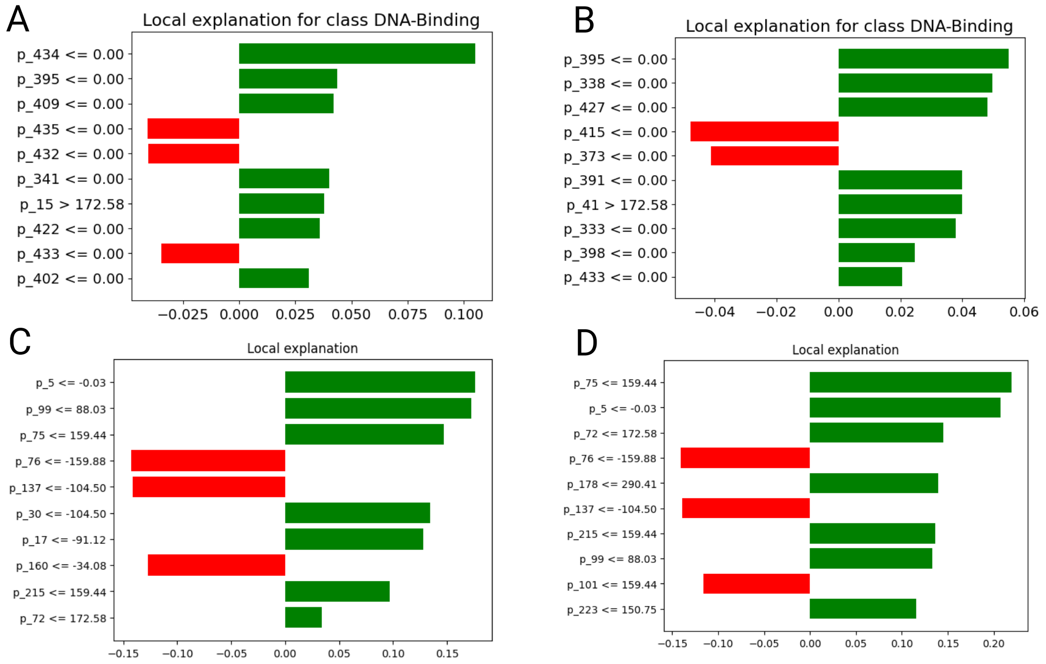

**Supplementary Figure S4**: **Local Interpretable Model-agnostic Explanations for classification and regression models explored in this work**. **A** Global summary of the DNA-binding classification model's main positions and physicochemical properties. **B** Global summary of the main positions and physicochemical properties for GFP regression model. **C, E, G** Single instances of SHAP values for DNA-Binding classification model. **D, F, H** Single instances of SHAP values for GFP regression model. **I** The ten most relevant positions and physicochemical values for the DNA-binding classification model. **J** The ten most relevant positions and physicochemical values for the GFP regression model.

In the context of the GFP predictive model, ICE plots and Partial Dependence Plots are generated to explore the most relevant properties identified through SHAP analysis. Figure **S5** illustrates how alterations in positions lead to corresponding changes in predictions, influencing the median brightness values either positively or negatively. The interpretation of the Partial Dependence Plots, depicted in Figure **S5**, reveals that modifying positions 82 and 67 has no discernible effect on the model's response when changes are negative. Conversely, positive changes correlate with an increase in the median brightness value. In essence, these plots serve as valuable tools for assisting in the design of protein sequences, enabling the exploration of prediction changes guided by sequence modifications. Moreover, this approach facilitates the identification of key residues crucial for maintaining specific functions or properties.

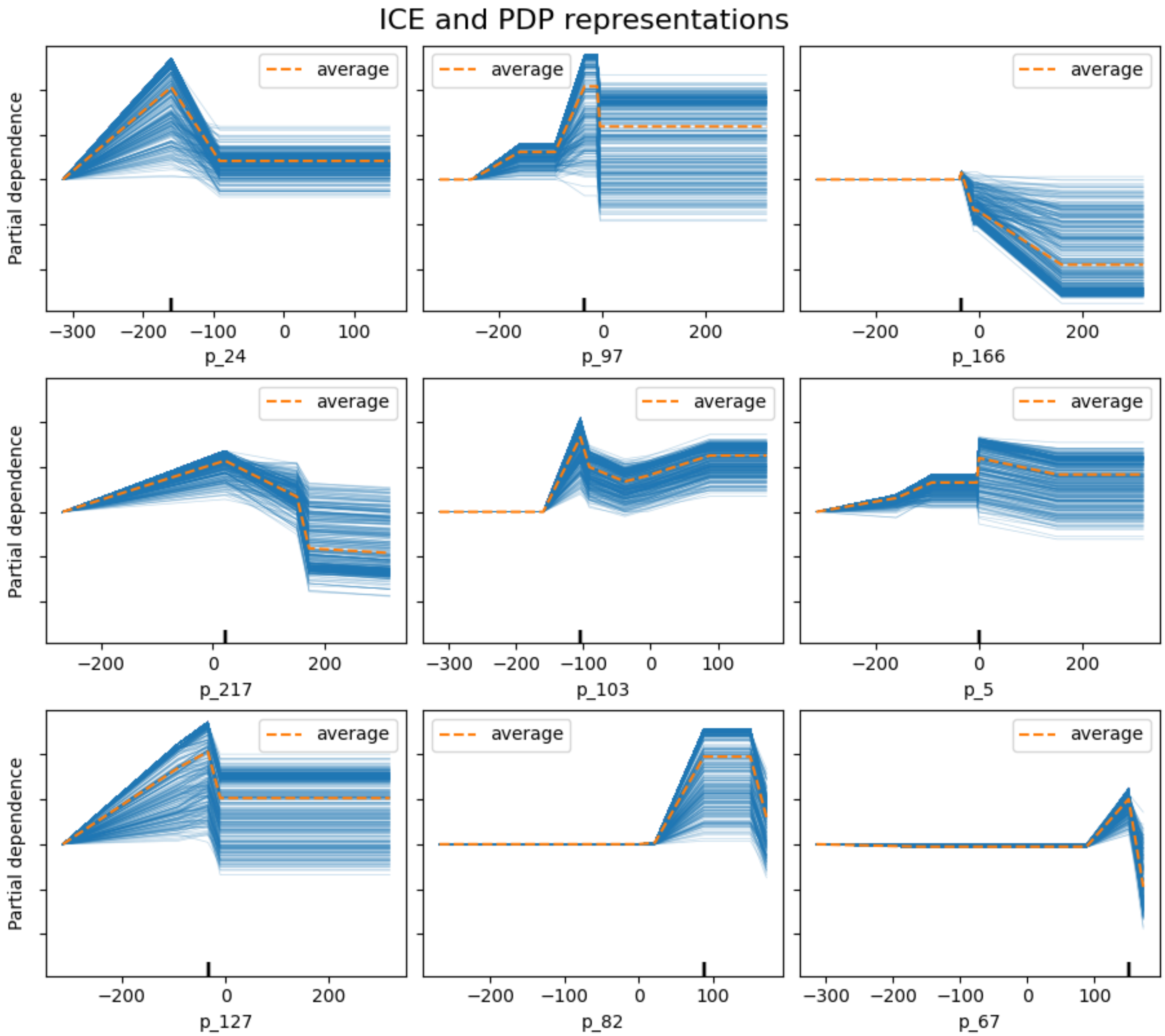

**Supplementary Figure S5**: **ICE plot and Partial dependence plot for the most relevant points identified by SHAP analysis for the GFP predictive model**. ICE plot visualization and partial dependence plot for the nine most relevant points identified by the SHAP analysis for the GFP predictive model. The X-axis show the relevant properties or descriptors, and the Y-axis represents the models' predictions. Changes in the property values affect the median brightness estimated positively or negatively. For example, changing the values in position 82 for positive values affects the model's predictions. The interpretation of this point is to introduce single-point mutations replacing the residues for arginine or aspartate.

Lundberg, S. M. and Lee, S.-I. (2017). A unified approach to interpreting model predictions. In Guyon, I., Luxburg, U. V., Bengio, S., Wallach, H., Fergus, R., Vishwanathan, S., and Garnett, R., editors, *Advances in Neural Information Processing Systems 30*, pages 4765–4774. Curran Associates, Inc.

Medina-Ortiz, D., Contreras, S., Amado-Hinojosa, J., Torres-Almonacid, J., Asenjo, J. A., Navarrete, M., and Olivera-Nappa, Á. (2022). Generalized property-based encoders and digital signal processing facilitate predictive tasks in protein engineering. *Frontiers in Molecular Biosciences*, 9.

Medina-Ortiz, D., Contreras, S., Quiroz, C., Asenjo, J. A., and Olivera-Nappa, Á. (2020a). Dmakit: A user-friendly web platform for bringing state-of-the-art data analysis techniques to non-specific users. *Information systems*, 93:101557.

Medina-Ortiz, D., Contreras, S., Quiroz, C., and Olivera-Nappa, Á. (2020b). Development of supervised learning predictive models for highly non-linear biological, biomedical, and general datasets. *Frontiers in molecular biosciences*, 7:13.

Mishra, A., Pokhrel, P., and Hoque, M. T. (2019). Stackdppred: a stacking based prediction of dna-binding protein from sequence. *Bioinformatics*, 35(3):433–441.

Mothilal, R. K., Sharma, A., and Tan, C. (2020). Explaining machine learning classifiers through diverse counterfactual explanations. In *Proceedings of the 2020 Conference on Fairness, Accountability, and Transparency*, pages 607–617.

Ribeiro, M. T., Singh, S., and Guestrin, C. (2016). "why should I trust you?": Explaining the predictions of any classifier. In *Proceedings of the 22nd ACM SIGKDD International Conference on Knowledge Discovery and Data Mining, San Francisco, CA, USA, August 13-17, 2016*, pages 1135–1144.

Sarkisyan, K. S., Bolotin, D. A., Meer, M. V., Usmanova, D. R., Mishin, A. S., Sharonov, G. V., Ivankov, D. N., Bozhanova, N. G., Baranov, M. S., Soylemez, O., et al. (2016). Local fitness landscape of the green fluorescent protein. *Nature*, 533(7603):397–401.

Siedhoff, N. E., Schwaneberg, U., and Davari, M. D. (2020). Machine learning-assisted enzyme engineering. *Methods Enzymol*, 643:281–315.
